# When *Vibrios* Take Flight: A Meta-analysis of Pathogenic *Vibrios* Species in Wild and Domestic Birds

**DOI:** 10.1101/2022.02.19.481111

**Authors:** Andrea J. Ayala, C. Brandon Ogbunugafor

**Affiliations:** Department of Ecology and Evolutionary Biology, Yale University, New Haven, Connecticut, 06520, USA

**Keywords:** Vibrio spp., Birds, Disease, Pathogenic, Ecology

## Abstract

Of the over 100 species in the genus *Vibrio*, approximately twelve are associated with clinical disease, such as cholera and vibriosis. Crucially, eleven of those twelve—*V. alginolyticus*, *V. cholerae*, *V. cincinnatiensis*, *V. hollinsae*, e.g., *Grimontia hollinsae*, *V. furnissii*, *V. mimicus*, *V. parahaemolyticus*, *V. vulnificus*, *V. harveyi*, *V. scophthalmi* and *V. metschnikovi*—have also been isolated from birds. Since 1965, pathogenic Vibrio species have been consistently isolated from aquatic and ground-foraging bird species, which has implications for public health, as well as the One Health paradigm defined as an ecology-inspired, integrative framework for the study of health and disease, inclusive of environmental, human, and animal health. In this meta-analysis, we identified 76 studies from the primary literature which report on or examine birds as hosts for pathogenic *Vibrio* species. We found that the burden of disease in birds was most commonly associated with *V. cholerae*, followed by *V. metschnikovi* and *V. parahaemolyticus*. Meta-analysis wide prevalences of the *Vibrio* pathogens varied from 19% for *V. parahaemolyticus* to 1% for *V. mimicus*. Wild and domestic birds were both affected, which may have implications for conservation, as well as agriculturally associated avian species. As pathogenic *Vibrios* become more abundant throughout the world as a result of warming estuaries and oceans, susceptible avian species should be continually monitored as potential reservoirs for these pathogens.

## 1 Introduction

Waterborne pathogens around the globe are experiencing a period of unprecedented global change, with the Vibrionaceae categorized among the most climate-sensitive families of aquatic prokaryotes [1, 2]. Evidence continues to mount concerning the uptick in the abundance, distribution, and phenology of the Vibrionaceae, since rising temperatures, humidity, and precipitation have led to their increased survival and rates of replication [3–5]. Within this family resides the genus Vibrio, a genetically diverse group of gram-negative, motile, and facultatively anaerobic bacteria that are endemic to marine and estuarine waters [6]. With over 100 named species in the Vibrio genus, approximately twelve are known to be pathogenic to human hosts [7]. Specifically, eleven of the twelve, i.e., *V. alginolyticus, V. cholerae, V. cincinnatiensis, V. hollinsae*, e.g., *Grimontia hollinsae, V. furnissii, V. mimicus, V. parahaemolyticus, V. vulnificus, V. harveyi, V. scophthalmi* and *V. metschnikovii*, are the causative agents of human vibriosis, a term that incorporates a broad range of clinical signs [8–10]. These pathogenic species arguably include some of the greatest public health burdens worldwide, and over the last 40 years, the incidence of *Vibrio* infections has strikingly increased [11–13]. The continued rise of the incidence and prevalence of *Vibrio* pathogens has contributed to an unprecedented worldwide health burden of enteric, diarrheal diseases [14, 15]. Yet, the Vibrionaceae are not only expanding their breadth throughout the human population – over the last one hundred and fifty years, it also appears that the *Vibrio* genus is expanding its niche into avian hosts, with ensuing implications for the One Health paradigm, and how we contextualize ‘human’ diseases [16–19].

During the fifth pandemic of cholera (1881-1886), the bacteriologist Gamelaia reported a disease afflicting Rock Pigeons (*Columba livia*) and domestic chickens (*Gallus gallus*) in southern Russia. It was described as “a disease of fowls”, of which the etiological agent was indistinguishable by morphological examination from *Vibrio cholerae* [20, 21]. This etiological agent would eventually be classified as *Vibrio metschnikovii*, and by the early twenty-first century, it would be considered one of twelve pathogenic *Vibrio* species that causes disease in human hosts [7, 22, 23]. The occurrence of another pathogenic Vibrio isolated from birds would not be reported until 1966, when individual species from the Gifu and Higashiyama Zoos (Table 2) in Japan tested positive by culture for Biotypes 1 and 2 of *Vibrio parahaemolyticus* [24]. Pathogenesis in these zoo birds was not reported [24]. Based on the literature, it is possible that the bird that had cultured positive for Biotype 2 of Vibrio parahaemolyticus was in fact shedding *V. alginolyticus* [25, 26].

Pathogenic *Vibrio* species can be lethal in human hosts. For example, *Vibrio vulnificus* is a causative agent of primary septicemia with a case fatality rate of up to fifty percent [27, 28]. As one of world’s leading causes of seafood-related deaths, *Vibrio vulnificus* is an opportunistic pathogen which causes high morbidity and mortality among the immunocompromised and those with liver disease [29, 30]. *Vibrio cholerae*, specifically serotypes O1 and O139, is likely the most well-known member of the *Vibrio* genus. It is a pathogen which has generated seven pandemics since 1817, and whose ecology and pathogenesis has been covered in depth [31–36]. *Vibrio parahaemolyticus* is a leading cause of seafood-borne illness, with clinicians reporting gastroenteritis and septicemia as the primary causes of morbidity among patients [37, 38]. *V. alginolyticus* and *V. fluvialis* are considered emerging pathogens and have been linked to gastroenteritis and extraintestinal infections [8, 39]. The remaining *Vibrio* species, *V. cincinnatiensis, V. hollinsae*, e.g., *Grimontia hollinsae, V. furnissii, V. mimicus, V. harveyi, V. scophthalmi* and *V. metschnikovii* have been linked to sporadic reports of disease in human hosts [40–45], however, that does not diminish their clinical, veterinary, or ecological importance.

The One Health paradigm is a collaborative endeavor that seeks to incorporate the health of the environment, animals and humans, given the understanding that the resilience of these individual components is integrated and intertwined [46, 47]. Thus, the emergence of pathogenic *Vibrio* species in birds is not only of public health importance [48, 49], but of significance to avian disease ecology, as little is known of the large-scale effects that members of the *Vibrio* genus may have upon species of conservation concern [50, 51]. With few exceptions [36], little is also known concerning the role that birds may play in the maintenance or potentially cyclical contamination of the brackish, aquatic reservoirs they share with other susceptible vertebrates [52–56]. Therefore, in this chapter, we build on the work of prior investigators who have identified the presence of pathogenic *Vibrio* species in avian species to a) identify the avian taxas most likely to excrete the pathogens and b) assess the prevalence of individuals in each community or sample that do so. We further examine whether pathogenic *Vibrio* species are immunogenic and/or pathogenic to birds, and the duration that they shed in experimental infection studies. We focus not just on studies that have identified the presence or absence of pathogenic *Vibrio* species in wild avian communities, but also include experimental infection and immunity studies. Our objective is to provide a baseline framework by which avian disease ecologists, wildlife management professionals, veterinarians, and One Health personnel can evaluate and/or mitigate the potential risks of emerging pathogenic *Vibrio* species within our wild birds.

## 2 Methods

Using Google Scholar and Web of Science [57, 58], we searched for peer-reviewed studies, pre-prints, abstracts, and graduate theses in which the antibodies against or the antigens of pathogenic *Vibrio* species were isolated from birds or from the avian environment (e.g. the isolation of *Vibrio* pathogens from avian fecal matter or their nests) [59]. In our search strategy, we used the following search terms and Boolean operators: “*Vibrio* pathogen of interest” OR “*Vibrio* pathogen and disease” and “bird*” OR “wild bird*” OR “avian” (n = 14,950). In our search, we systematically searched for studies that examined evidence of infection by the following members of the *Vibrio* genus: *V. alginolyticus, V. cholerae, V. cincinnatiensis, V. furnissii, V. mimicus, V. parahaemolyticus, V. vulnificus, V. harveyi, V. scophthalmi* and *V. metschnikovii*. Given the relatively recent taxonomic reclassifications of *Grimontia hollinsae* [60] and *Photobacterium damselae* [61] from the genus *Vibrio*, we also included these pathogens in our analysis. We included experimental infection studies, case reports, and cross-sectional studies published between 1966 and January 1, 2022. In our analysis, we excluded sources that did not serve as primary literature involving investigations of pathogenic *Vibrio* infections in domestic or wild birds, e.g., retrospective studies and review papers (retrospective and review papers, n = 24) as well as duplicates (duplicates, n = 81). We also excluded any literature without a clear diagnostic and physiological association between domestic or wild birds as hosts of our *Vibrio* species of interest (exclusion criteria, n = 14845).

From each study, we extracted the following elements when available: avian species or taxonomic grouping, *Vibrio* species, country, year the study was conducted or published, the number of birds tested or infected, the number of birds from which *Vibrio* was isolated, and the method(s) by which pathogenic *Vibrio* species were identified. Where possible, we identified the prevalence of our *Vibrio* pathogens of interest, including presence and absence, to determine study-wide prevalence. We also reported serotypes and/or clinically important strains when that information was provided. We further identified whether our *Vibrio* pathogens of interest were associated with clinical signs or avian mortality events, however, unless specifically stated in the text, we could not determine whether our *Vibrio* pathogens of interest were the causative agent(s) of reported morbidity or mortality.

## 3 Results

### 3.1 Literature Review

We identified 76 studies from the primary literature that met our inclusion criteria, resulting in 425 study records of avian species or taxonomic groups from which the presence or absence of pathogenic *Vibrio* species was recorded (identified species, n = 171, identified families, n = 46). In our meta-analysis, a study record ranges from a single examined bird to 565 examined birds, which reflects the same species or taxonomic group that was tested for a single pathogenic *Vibrio* species of interest that originated from within the same study (Tables 1-5). In our meta-analysis, 29 countries were represented, constituting all continents except for Antarctica. Of the fifty-five years between 1966 and the start of 2022, studies were either published in or conducted in 41 of them. Sixteen study records did not provide sufficient information from which to identify *Vibrio* prevalence, as either the number of birds, flocks, nests, or sites were incompletely reported, or collected samples were pooled. When the *Vibrio* pathogens of interest were either not named or not classified to species, it was categorized as “*Vibrio* spp.”.

**Table 1:**
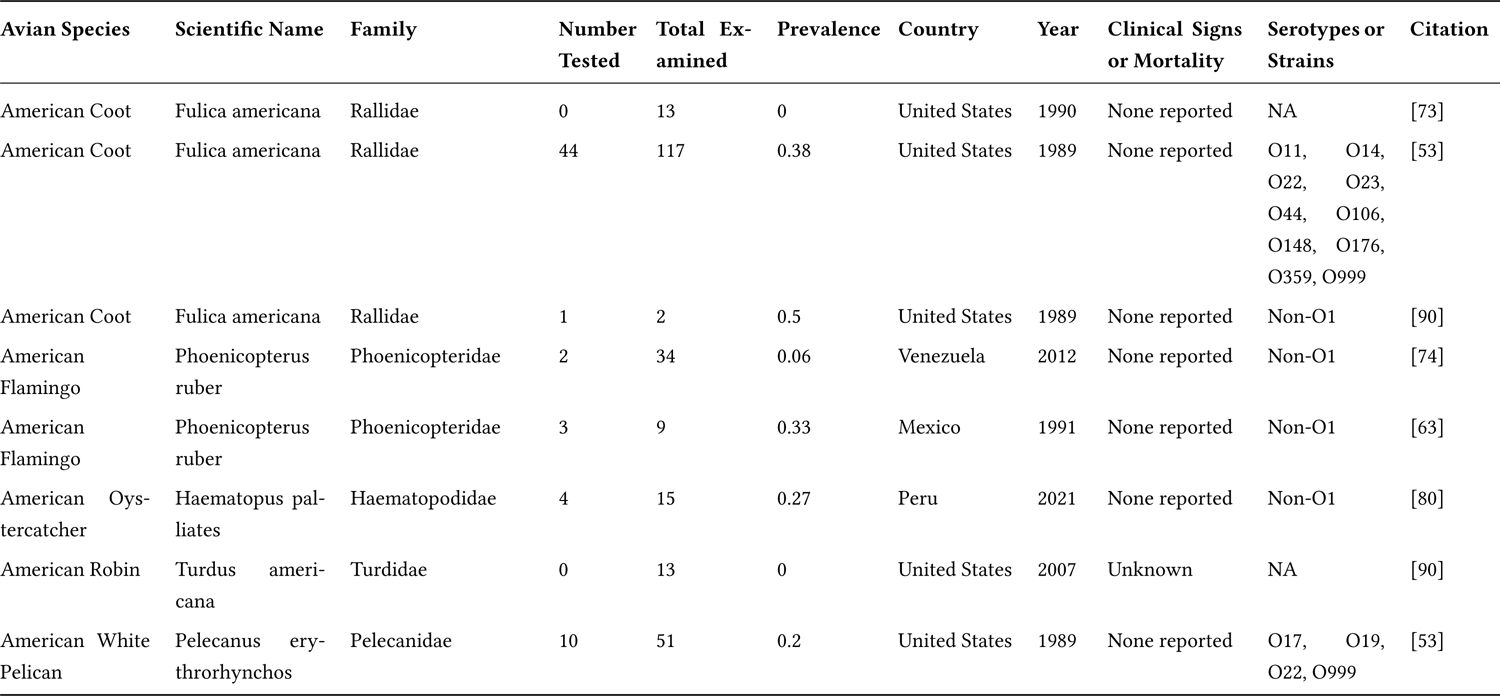
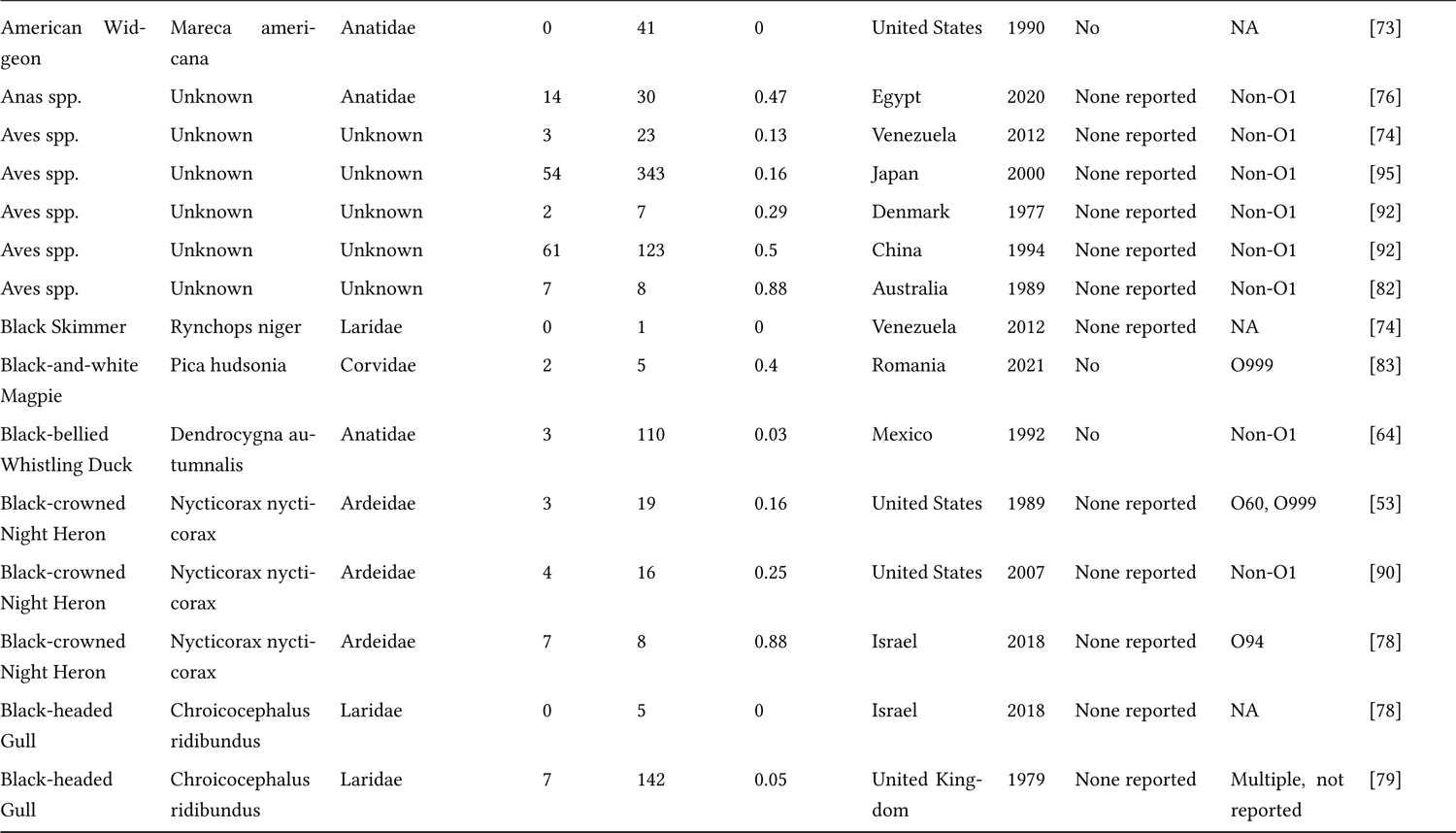
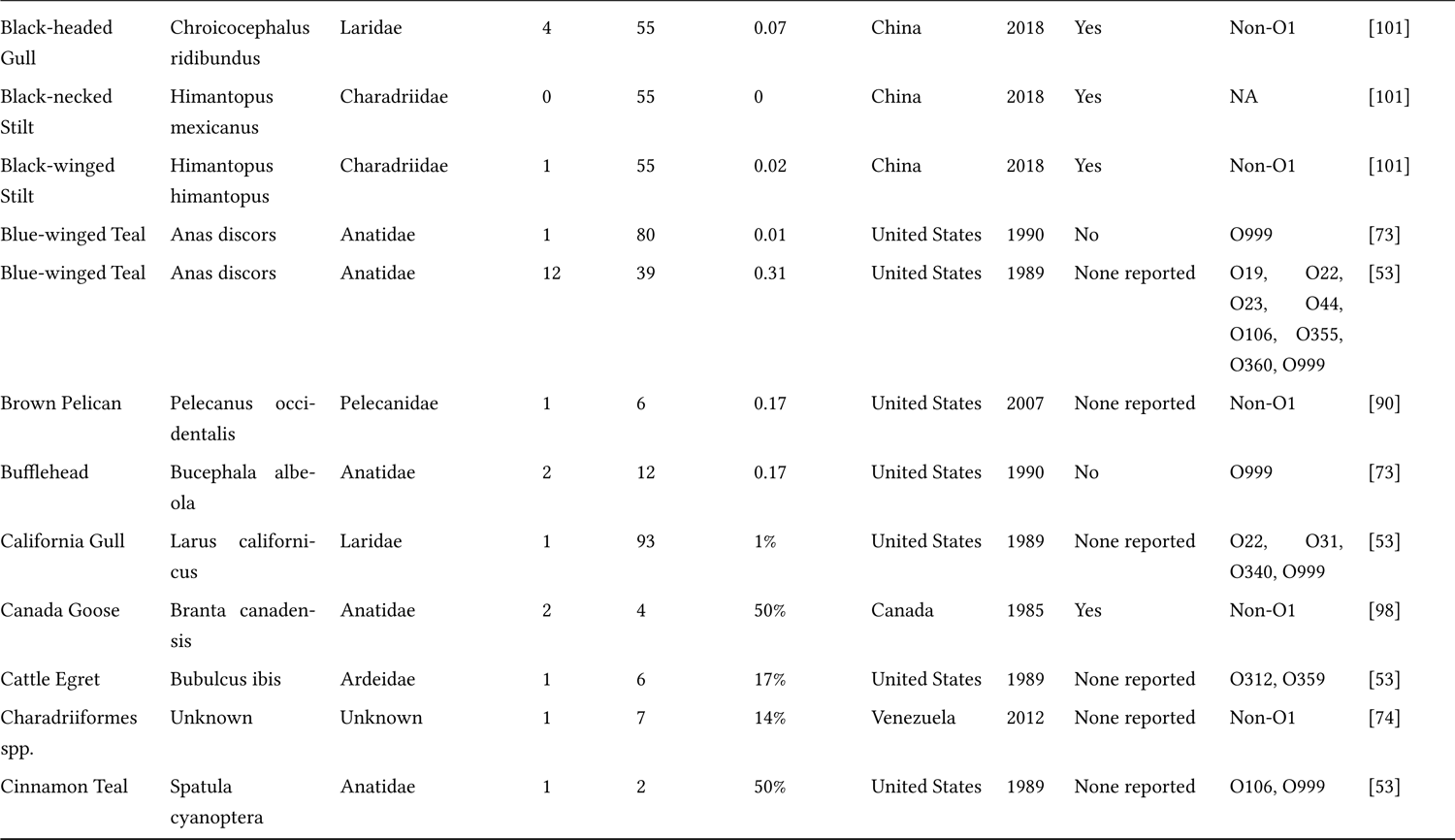
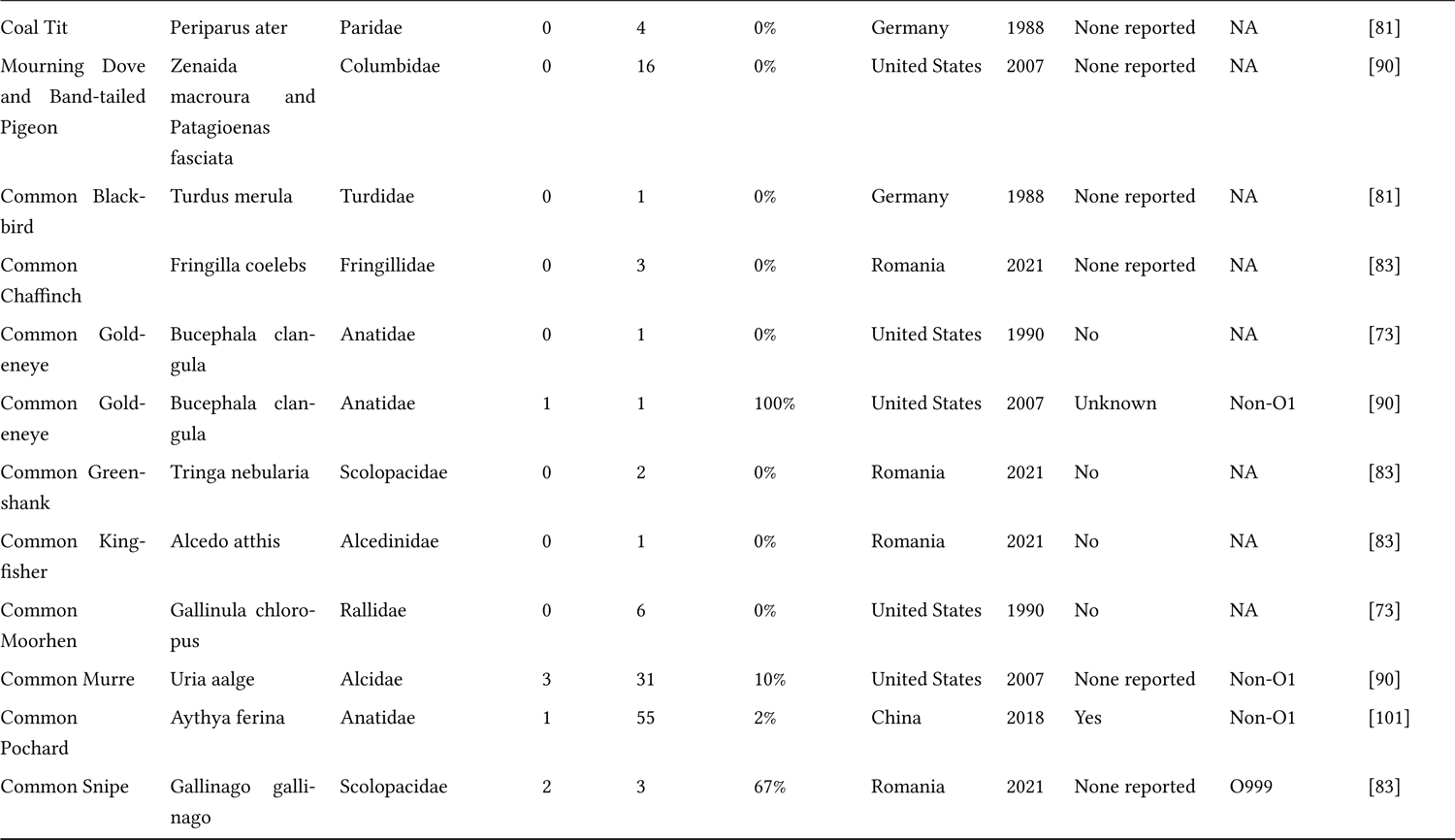
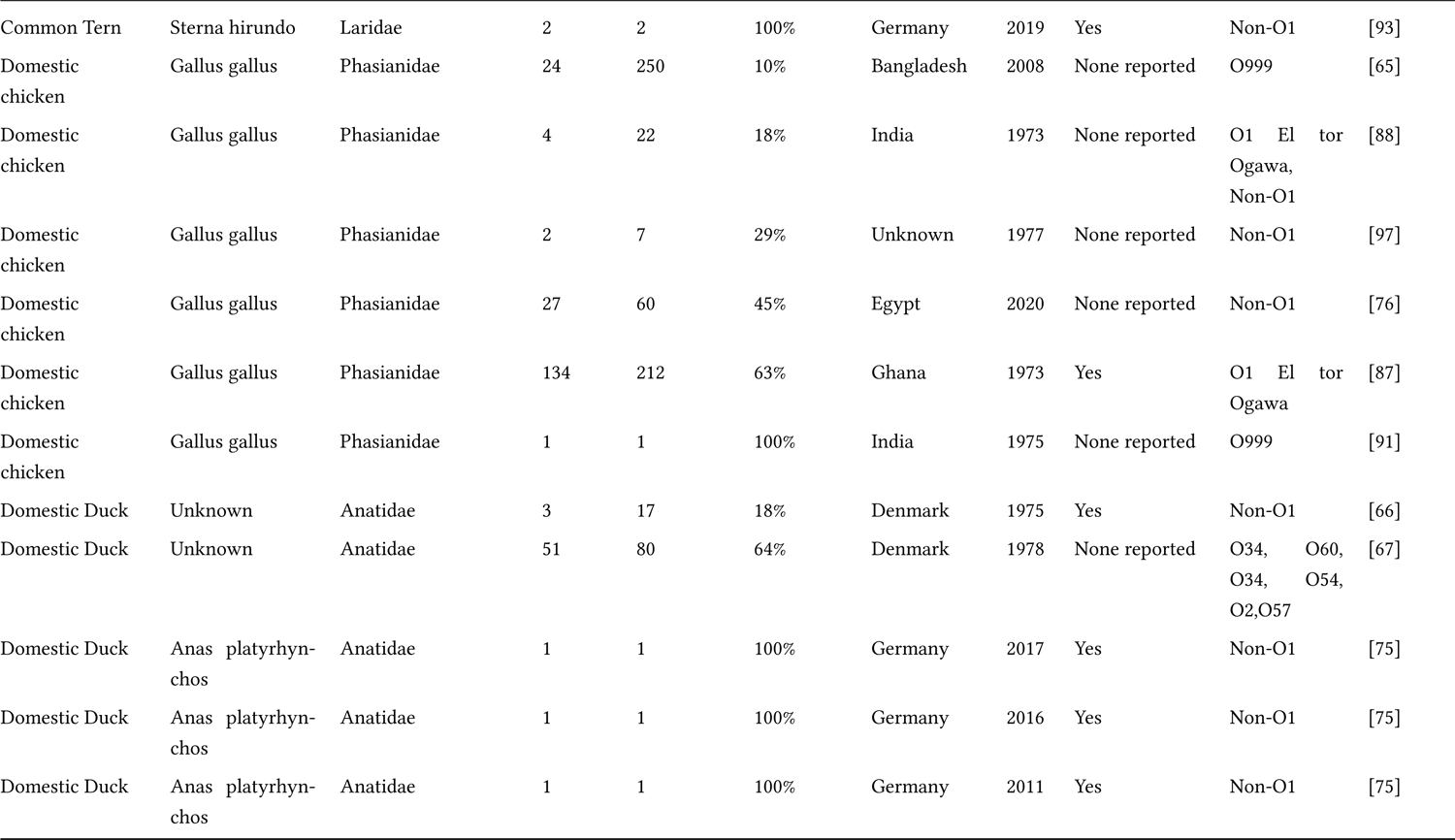
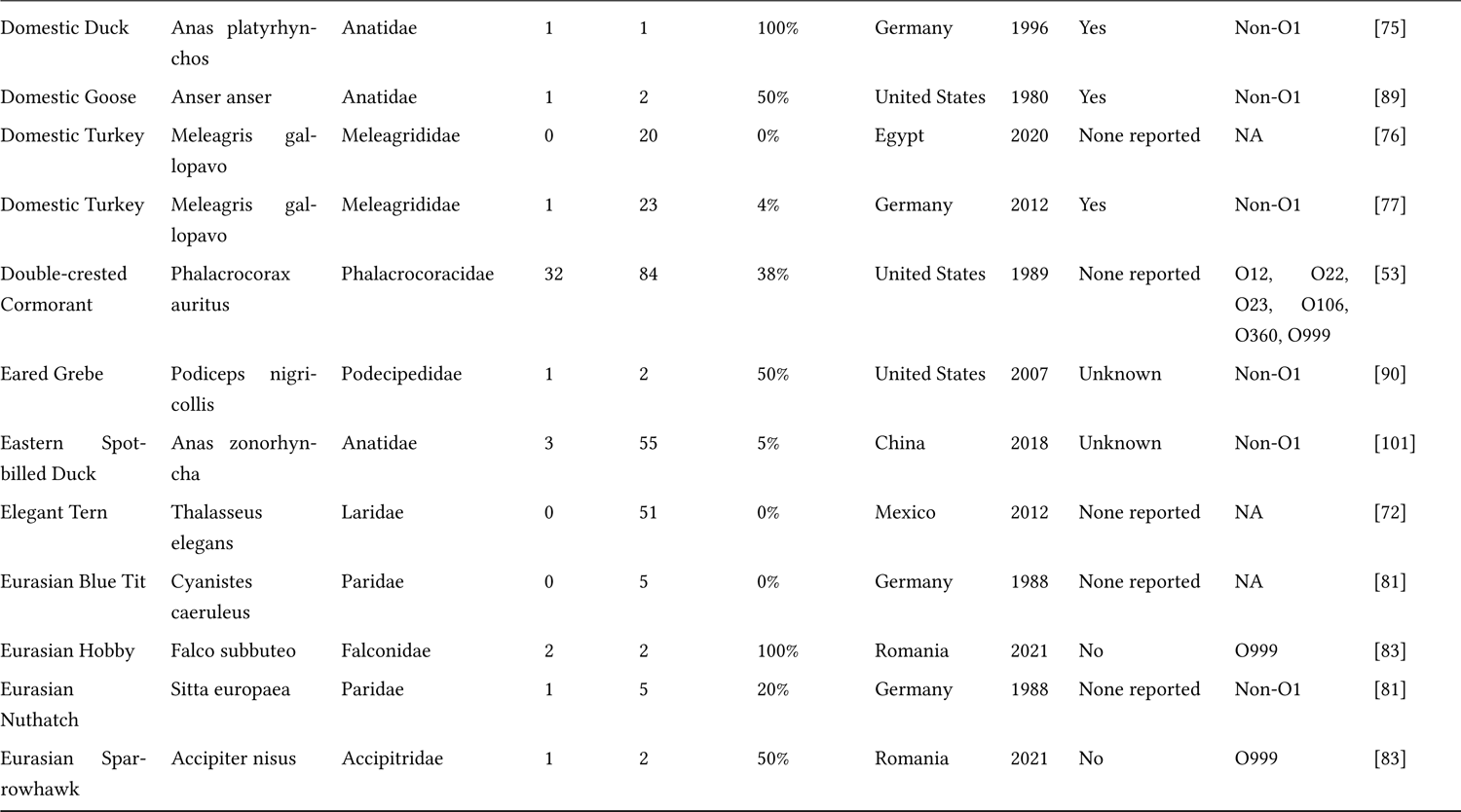
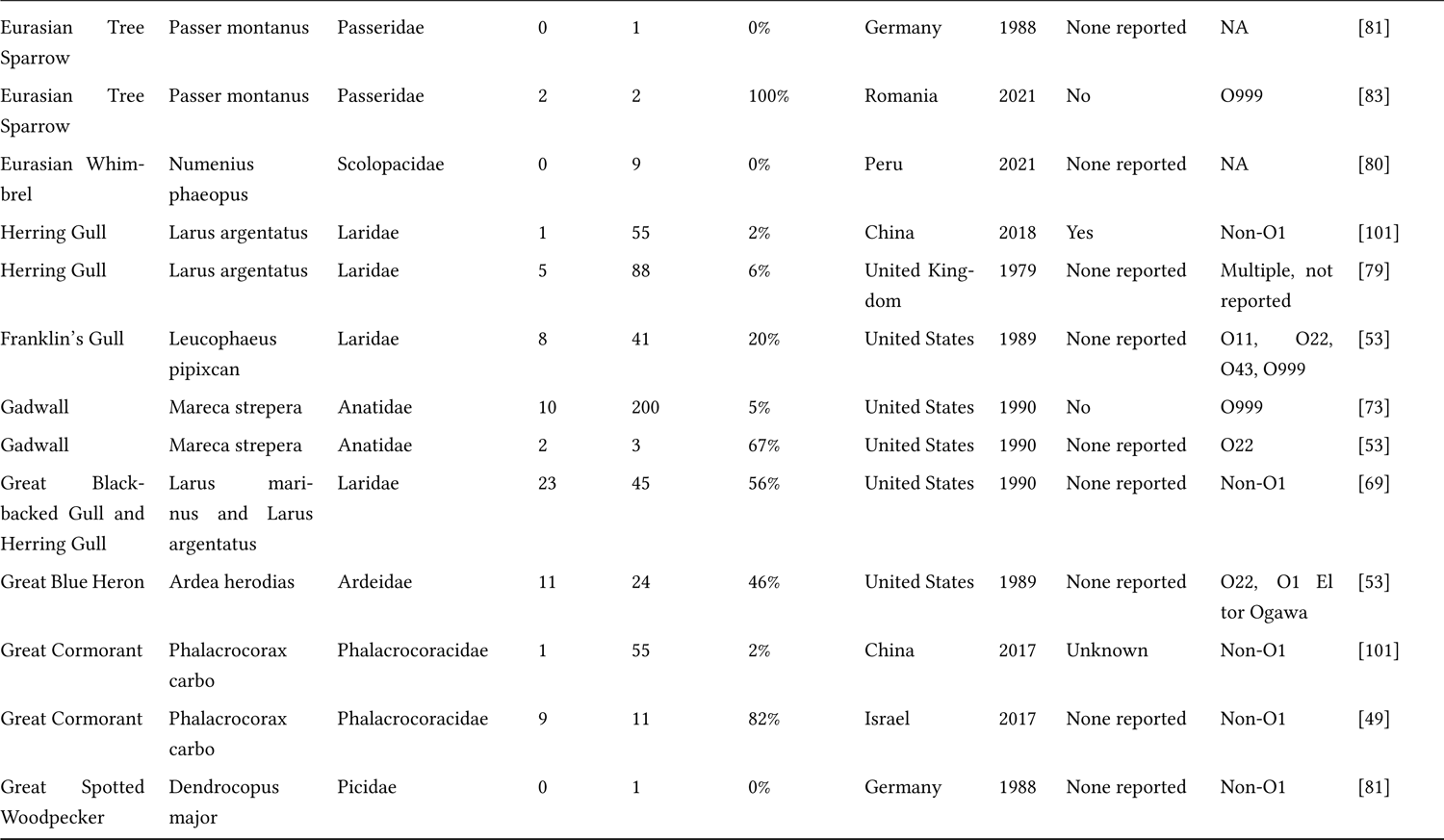
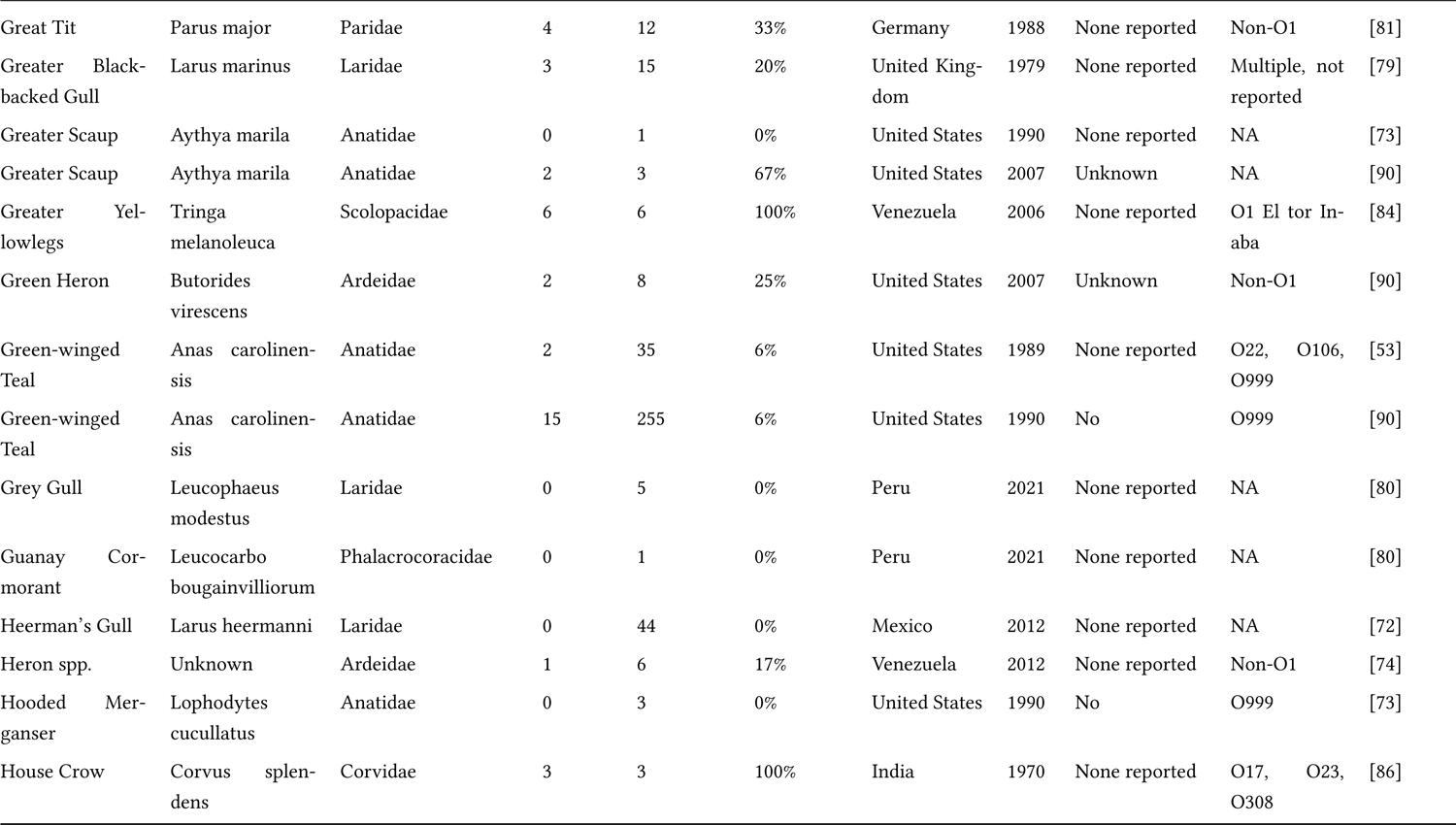
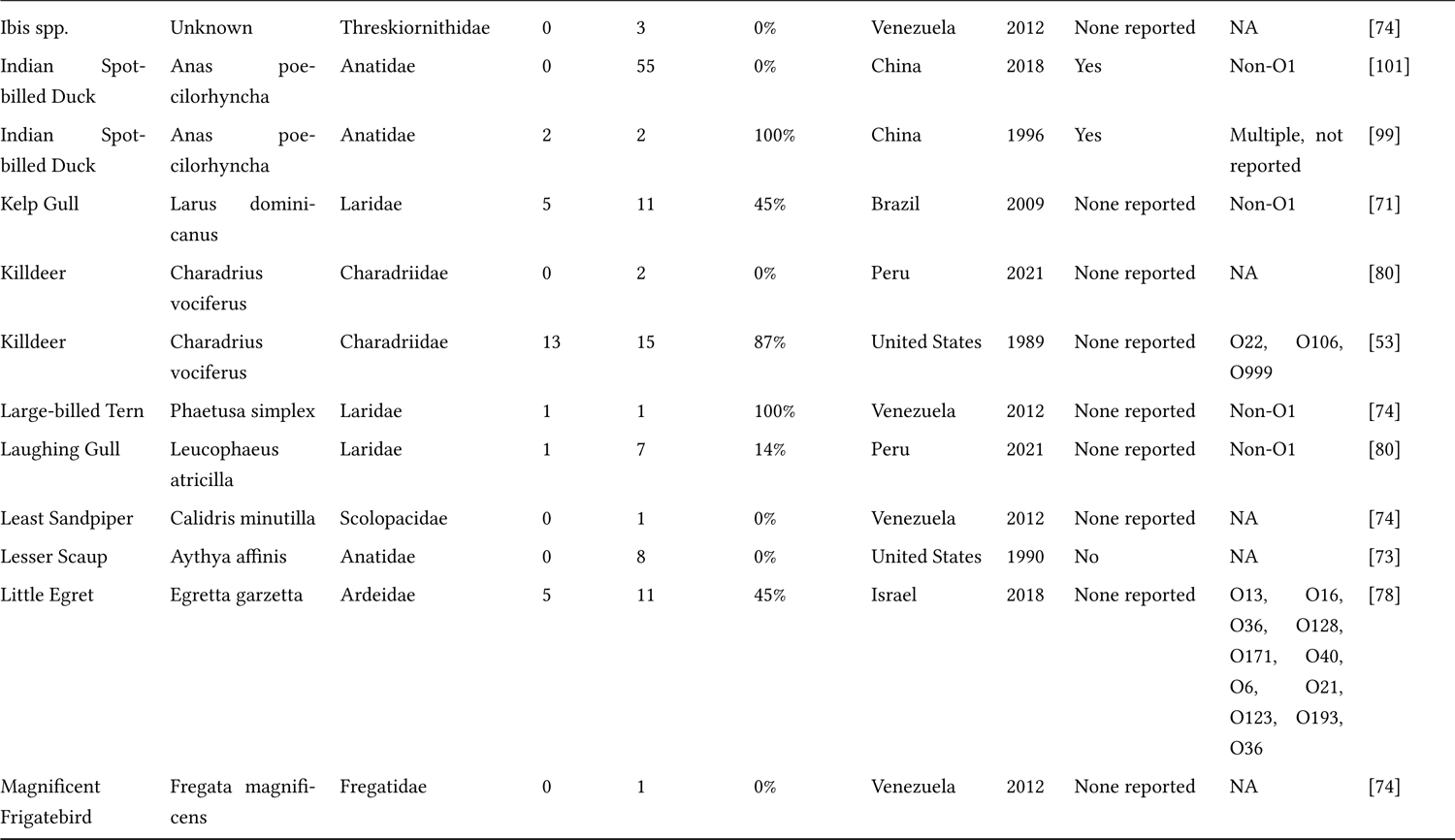
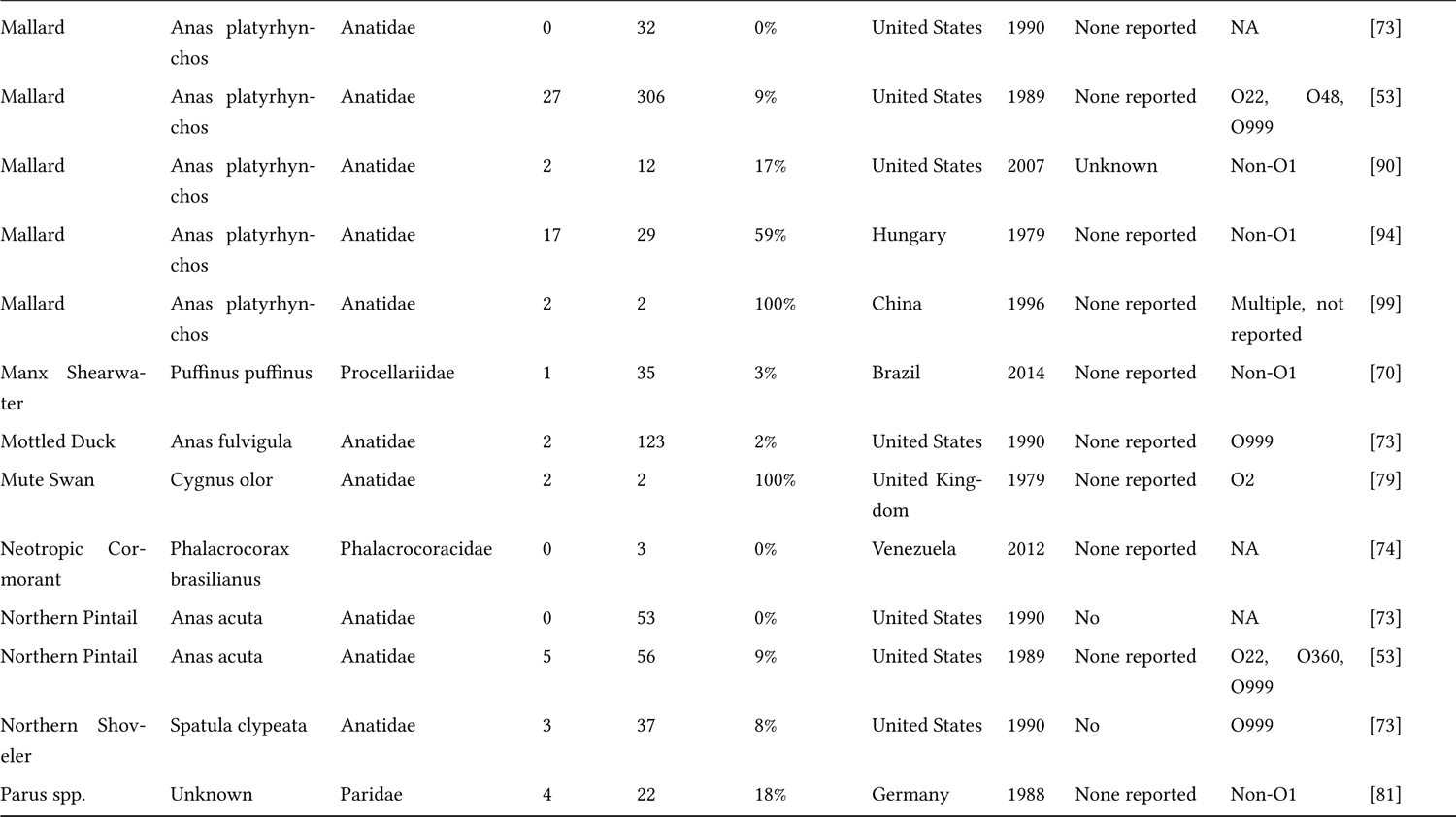
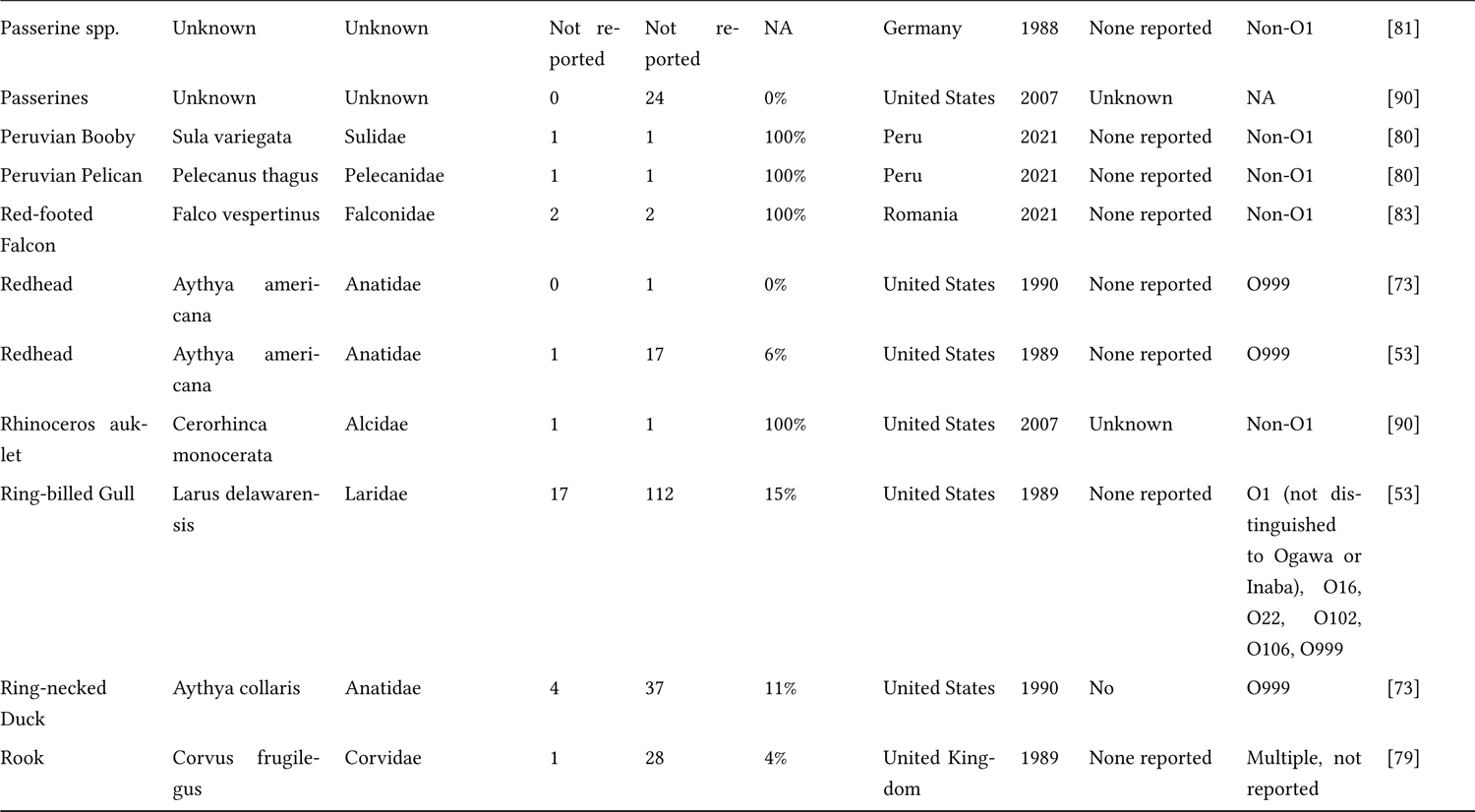
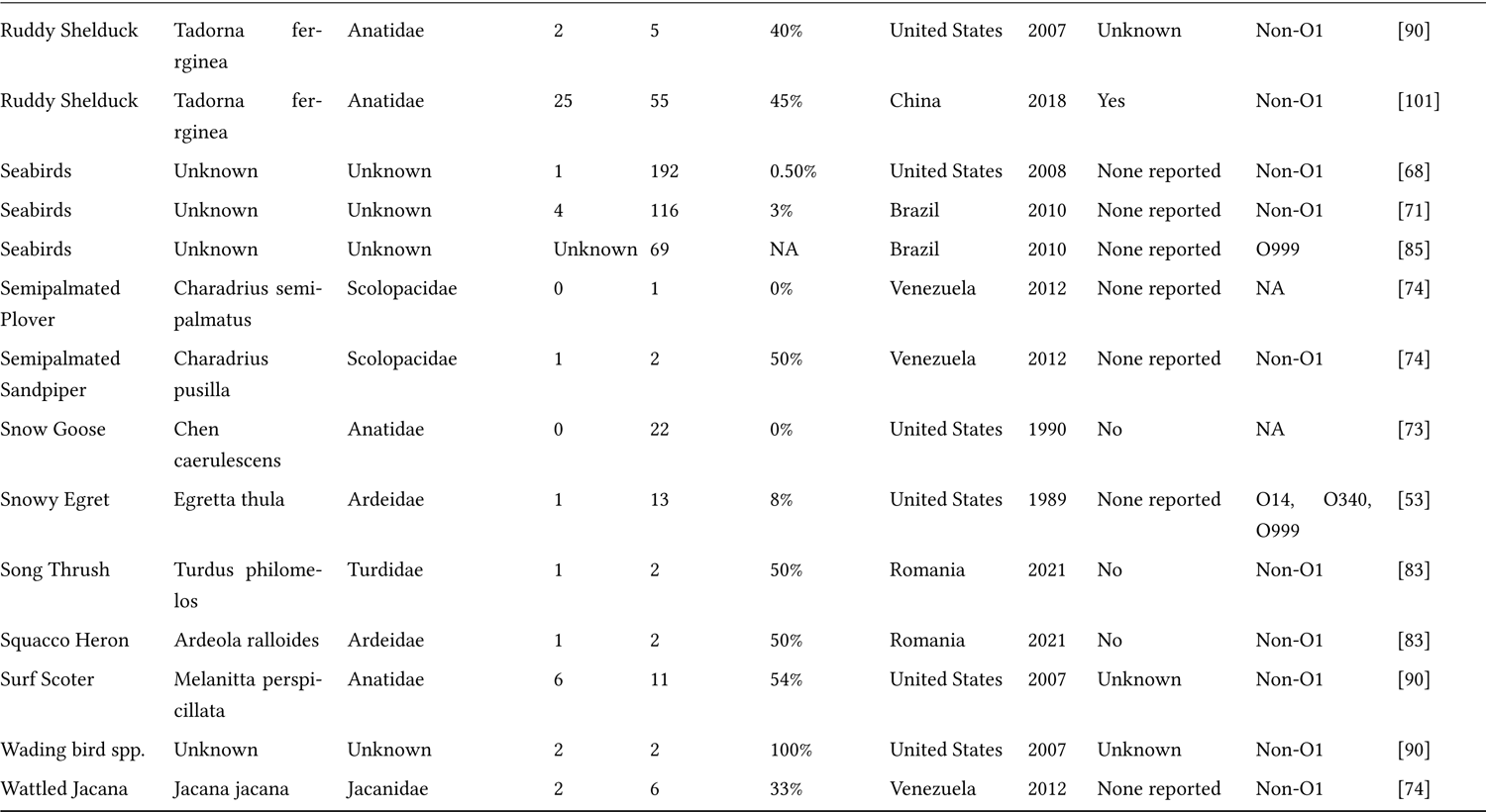
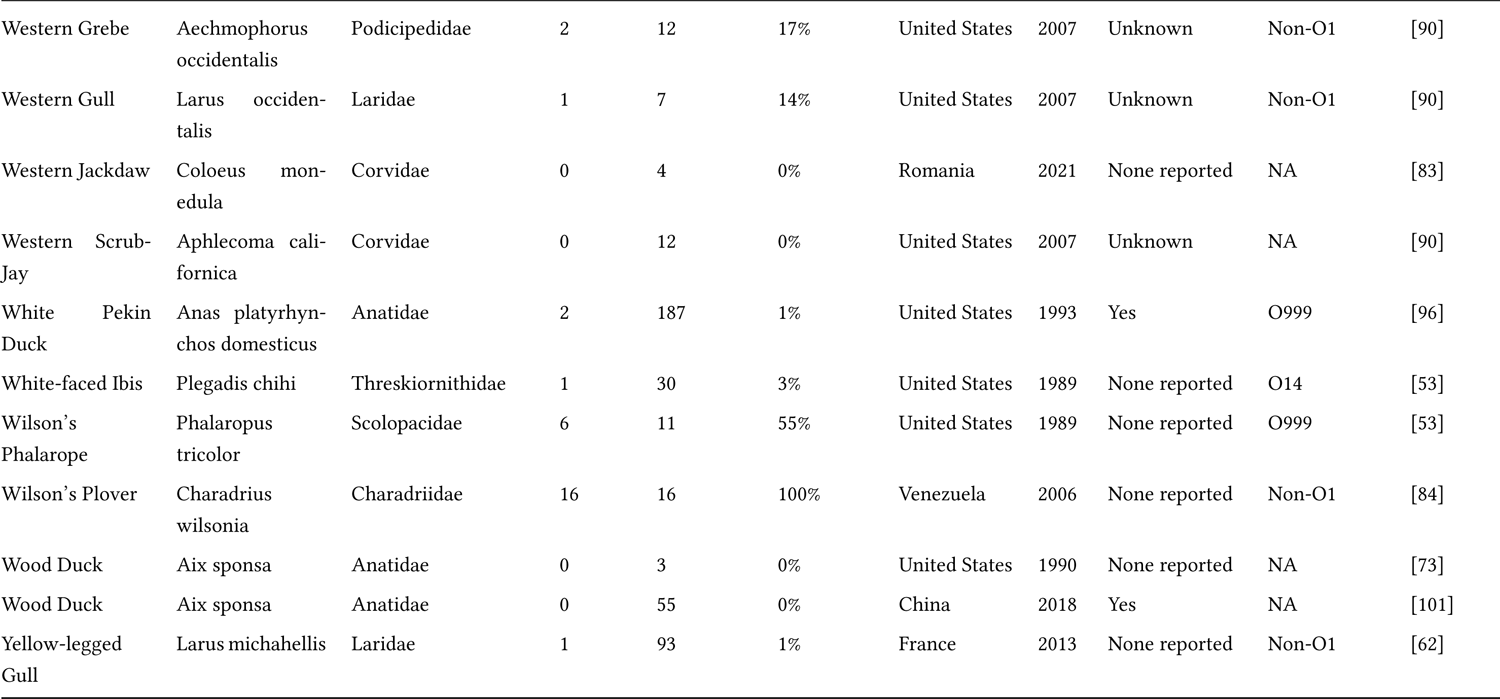
This table details the study records extracted from the 41 studies that investigated birds as hosts for *Vibrio cholerae*, both O1 El tor and Non-O1. In the table are provided the common name, the scientific name, the family, the number of birds that tested positive for *Vibrio cholerae*, the total number of birds examined for *V. cholerae*, and the prevalence for that record. We also provide the country that the study was performed in, as well as the year the study was conducted or published. For studies that reported clinical signs or mortality, we placed a “Yes” in that column. For serotypes or strains, there were several primary categories, for Non-O1, this involves any *Vibrio cholerae* that was tested for the properties specific to *V. cholerae* O1. The designation O999 was used when no typing was utilized beyond species, e.g., the serotype of *V. cholerae* was not reported or investigated further. NA was provided when the prevalence was zero, and no serotype was applicable. When multiple serotypes were reported, but not tied to records, we specified that in the column, and when serotypes were provided and could be tied to a record, we noted that in the column as well.

**Table 2:**
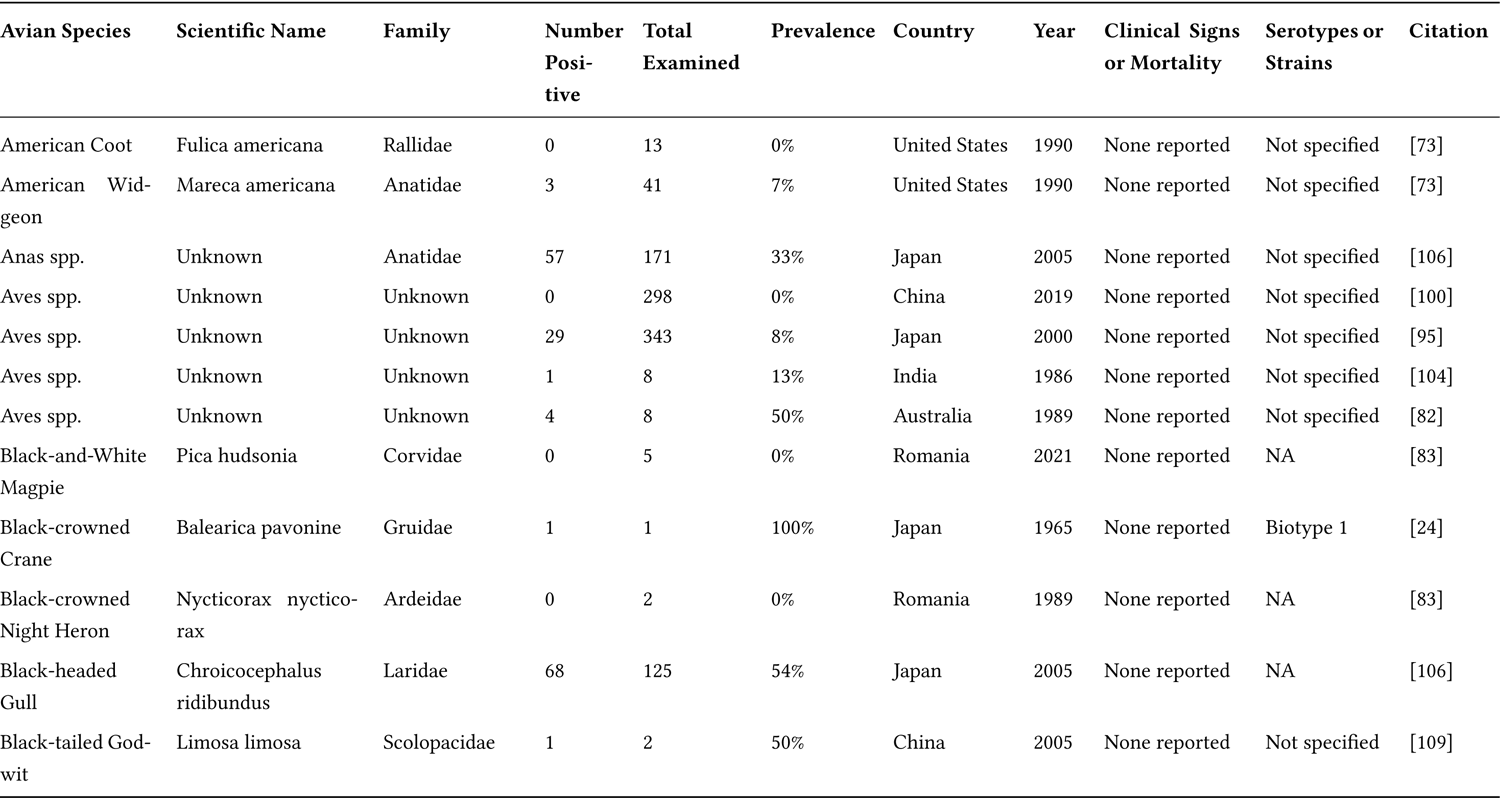
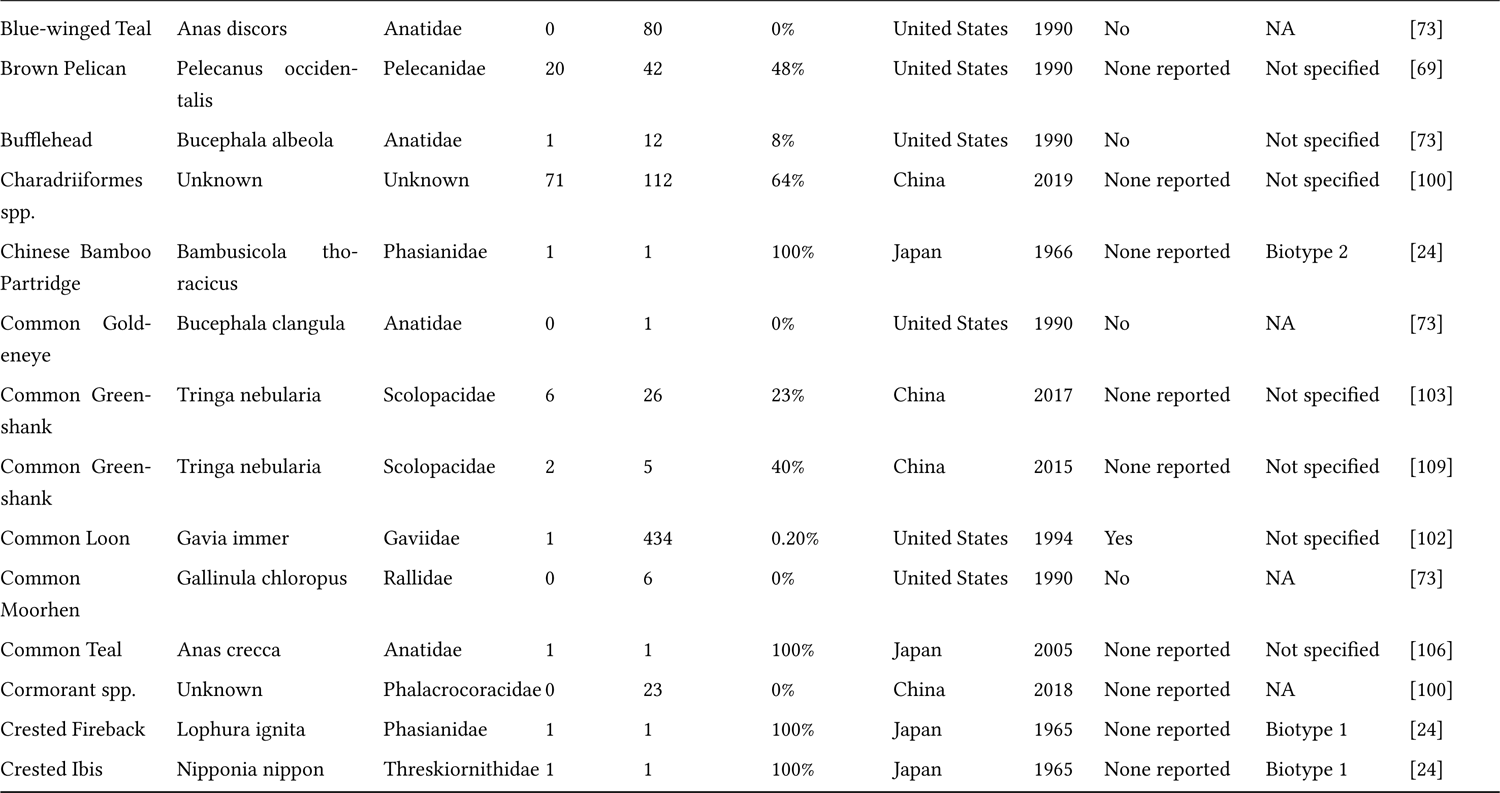
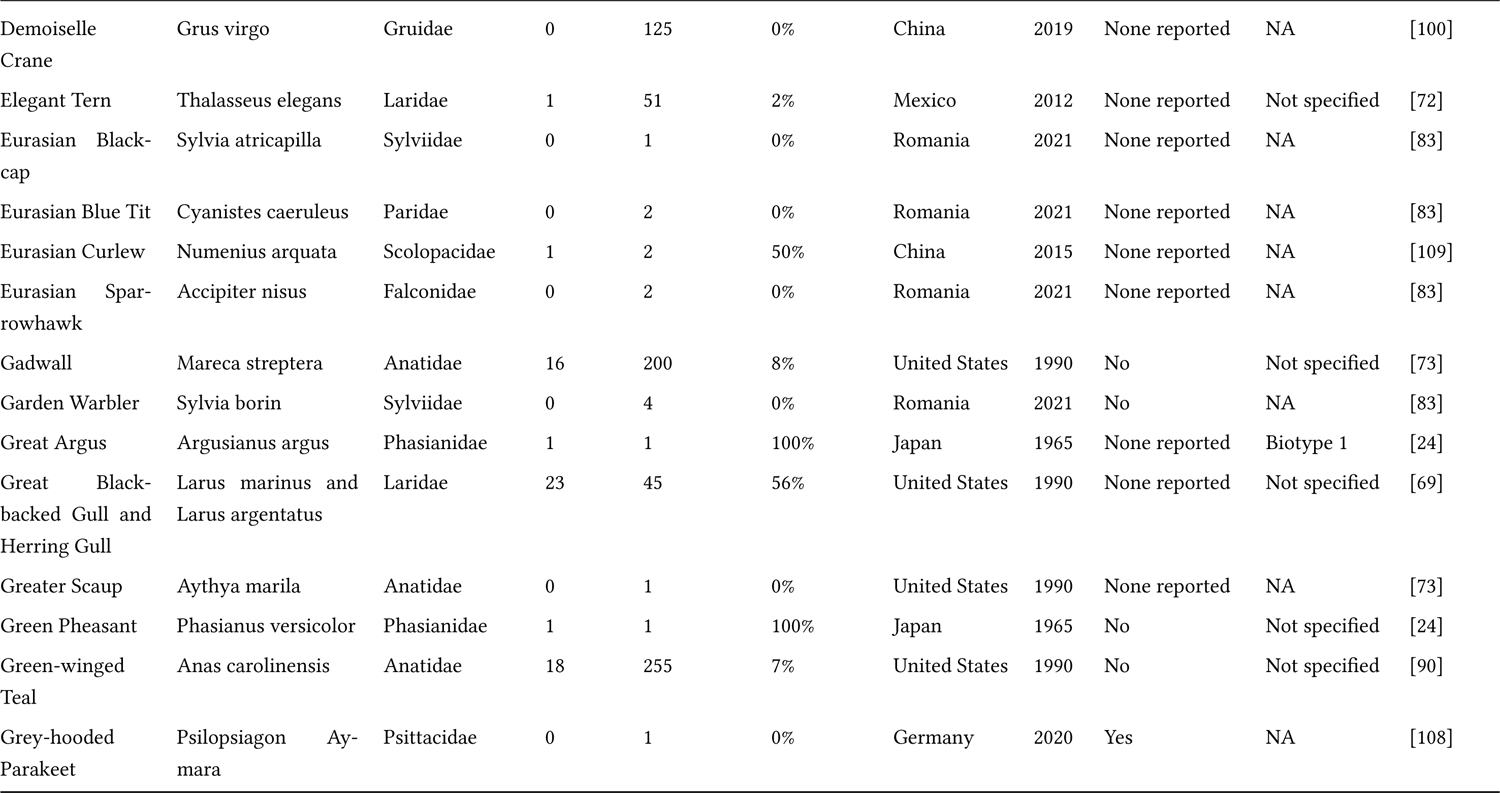
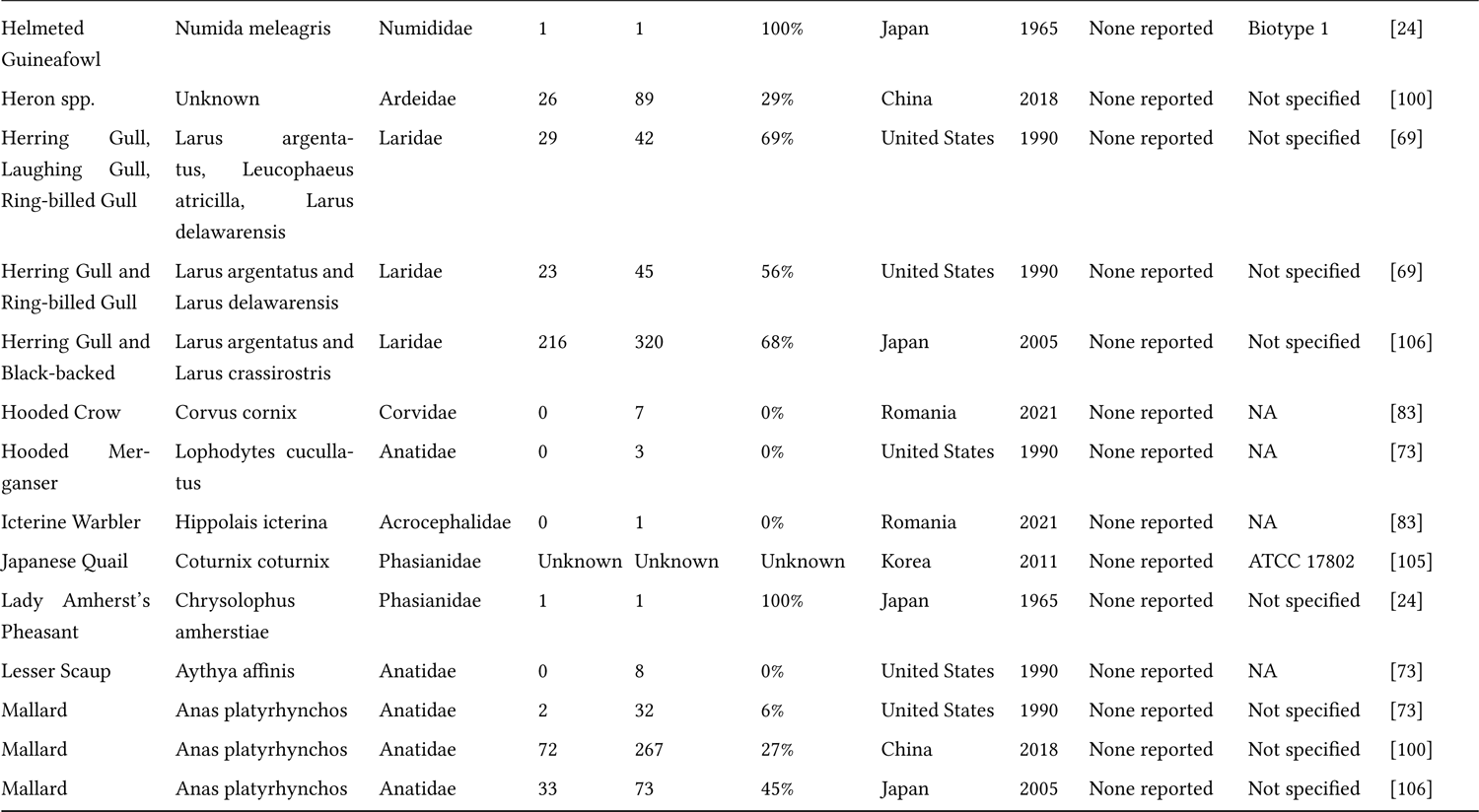
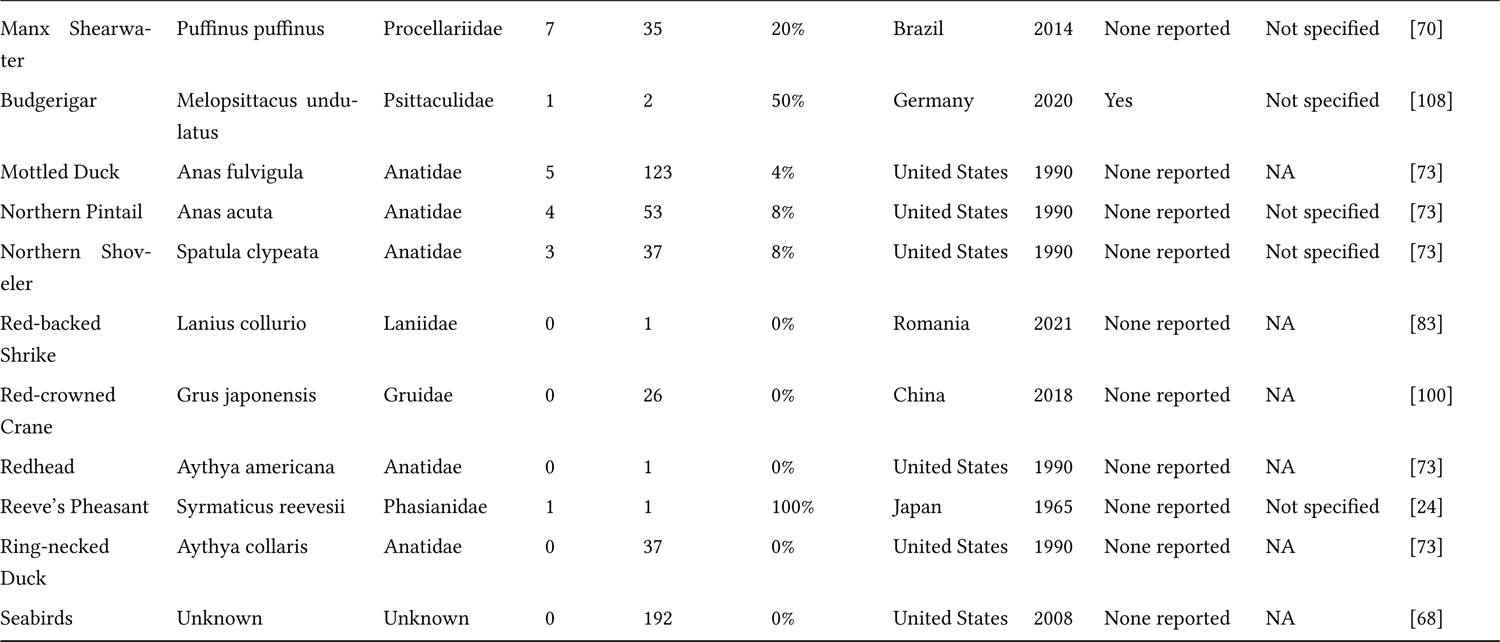
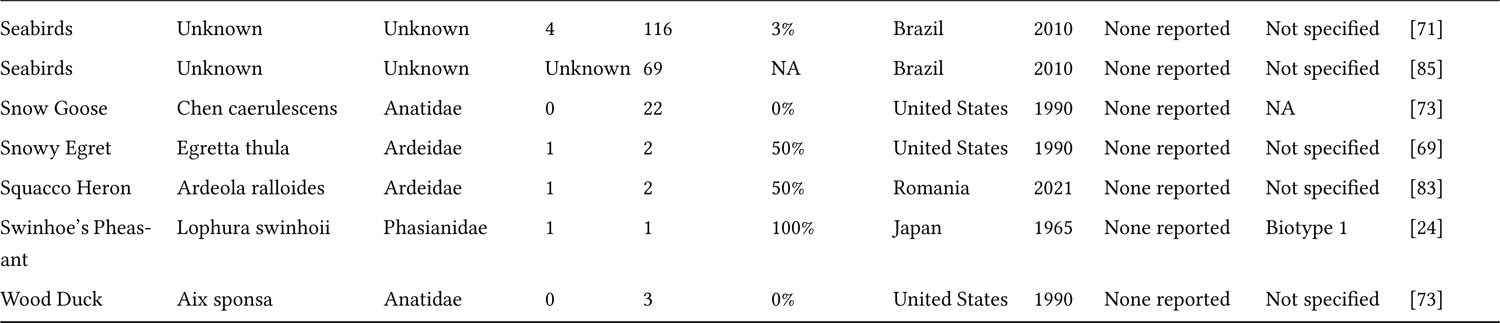
This table details the study records extracted from the 20 studies that investigated birds as hosts for *Vibrio parahaemolyticus*. In the table are provided the common name, the scientific name, the family, the number of birds that tested positive for *Vibrio parahaemolyticus*, the total number of birds examined for *V. parahaemolyticus*, and the prevalence for that record. We also provide the country that the study was performed in, as well as the year the study was conducted or published. For studies that reported clinical signs or mortality, we placed a “Yes” in that column. For serotypes or strains, we identified the strain when it was available. NA was provided when the prevalence was zero, and no strain was applicable.

**Table 3:** This table details the records extracted from the eight studies that investigated birds as hosts for Vibrio vulnificus. In the table are provided the common name, the scientific name, the family, the number of birds that tested positive for *Vibrio vulnificus*, the total number of birds examined for the pathogen, and the prevalence for that record. We also provide the country that the study was performed in, as well as the year the study was conducted or published. For studies that reported clinical signs or mortality, we placed a “Yes” in that column. For serotypes or strains that were reported, we reported those as well, otherwise we designated the column as not specified.

| Avian Species | Scientific Name | Family | Tested Number | Total Examined | Prevalence | Country | Year | Clinical Signs or Mortality | Serotypes or Strains | Citation |
| --- | --- | --- | --- | --- | --- | --- | --- | --- | --- | --- |
| American Crow | Corvus brachyrhynchos | Corvidae | Unknown | Unknown | Unknown | United States | 2019 | None reported | NA | [111] |
| Barred Warbler | Curruca nisoria | Sylviidae | 0 | 3 | 0% | Romania | 2021 | None reported | NA | [83] |
| Black-crowned Night Heron | Nycticorax nycticorax | Ardeidae | 0 | 2 | 0% | Romania | 2021 | None reported | NA | [83] |
| Black-headed Gull | Chroicocephalus ridibundus | Laridae | 1 | 125 | 1% | Japan | 2005 | None reported | Not specified | [106] |
| Black-tailed Godwit | Limosa limosa | Scolopacidae | 1 | 2 | 50% | China | 2015 | None reported | Not specified | [109] |
| Common Green-shank | Tringa nebularia | Scolopacidae | 0 | 5 | 0% | China | 2015 | None reported | Not specified | [109] |
| Common Teal | Anas crecca | Anatidae | 0 | 1 | 0% | Japan | 2005 | None reported | NA | [106] |
| Domestic chicken | Gallus gallus | Phasianidae | 4 | 565 | 0.70% | Nigeria | 2017 | None reported | Not specified | [110] |
| Eurasian Blue Tit | Cyanistes caeruleus | Paridae | 0 | 2 | 0% | Romania | 2021 | None reported | NA | [83] |
| Common Teal | Anas crecca | Anatidae | 0 | 2 | 0% | Romania | 2021 | None reported | NA | [83] |
| Garden Warbler | Sylvia borin | Sylviidae | 0 | 4 | 0% | Romania | 2021 | None reported | NA | [83] |
| Herring Gull and Black-tailed Gull | Larus argentatus and Larus crassirostris | Laridae | 86 | 320 | 27% | Japan | 2005 | None reported | NA | [106] |
| Hooded Crow | Corvus cornix | Corvidae | 0 | 7 | 0% | Romania | 2021 | None reported | NA | [83] |
| Japanese Quail | Coturnix coturnix | Phasianidae | Unknown | Unknown | Unknown | Korea | 2011 | None reported | KCTC 2959 | [105] |
| Laughing Gull | Leucophaeus atricilla | Laridae | Unknown | Unknown | Unknown | United States | 2021 | None reported | Not specified | [111] |
| Lesser Whitethroat | Sylvia curruca | Sylviidae | 0 | 2 | 0% | Romania | 2021 | Not reported | NA | [83] |
| Mallard | Anas platyrhynchos | Anatidae | 0 | 73 | 0% | Japan | 2005 | None reported | NA | [106] |
| Muscovy Duck | Cairina moschata | Anatidae | Unknown | Unknown | Unknown | United States | 2019 | None reported | Not specified | [111] |
| Seabirds | Unknown | Unknown | 2 | 116 | 2% | Brazil | 2010 | None reported | Not specified | [71] |
| Seabirds | Unknown | Unknown | Unknown | 69 | NA | Brazil | 2010 | None reported | Not specified | [85] |
| Wood Sandpiper | Tringa glareola | Scolopacidae | 0 | 2 | 0% | Romania | 2021 | Not reported | NA | [83] |

**Table 4:**
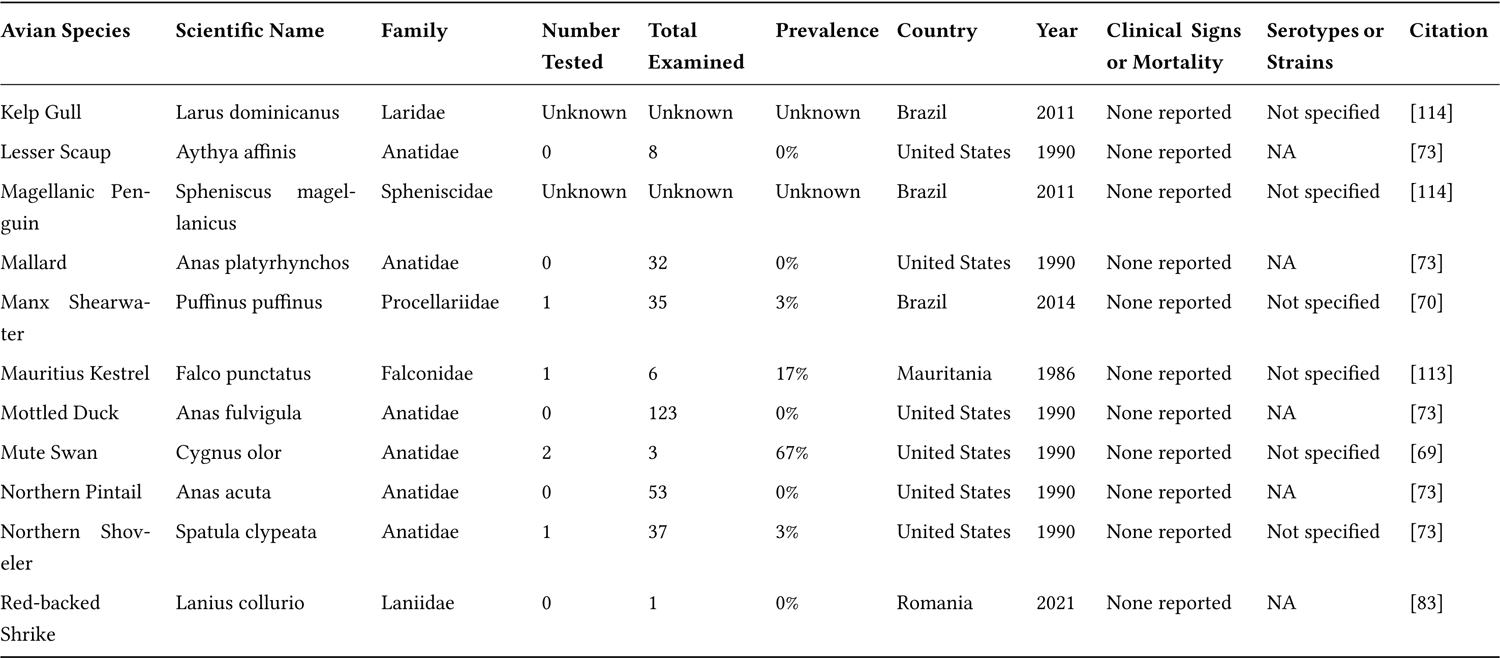
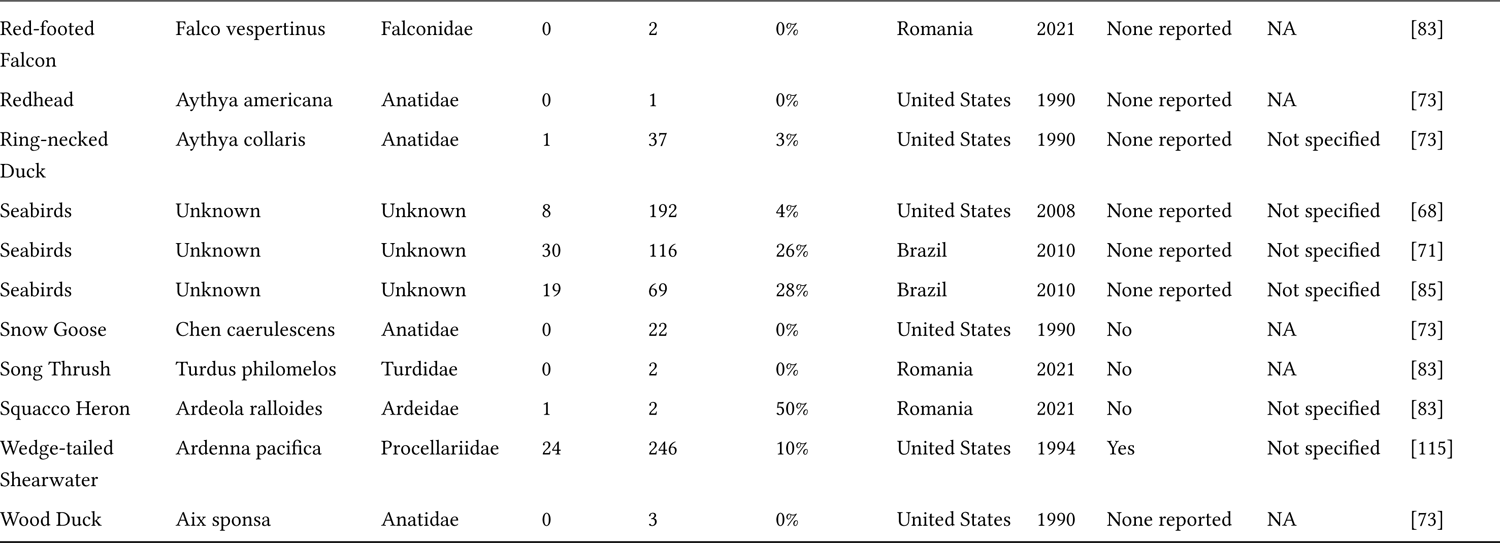
This table details the records extracted from the 15 studies that investigated birds as hosts for *Vibrio alginolyticus*. In the table are provided the common name, the scientific name, the family, the number of birds that tested positive for *Vibrio cholerae*, the total number of birds examined for *Vibrio alginolyticus*, and the prevalence for that record. We also provide the country that the study was performed in, as well as the year the study was conducted or published. For studies that reported clinical signs or mortality, we placed a “Yes” in that column.

**Table 5:**
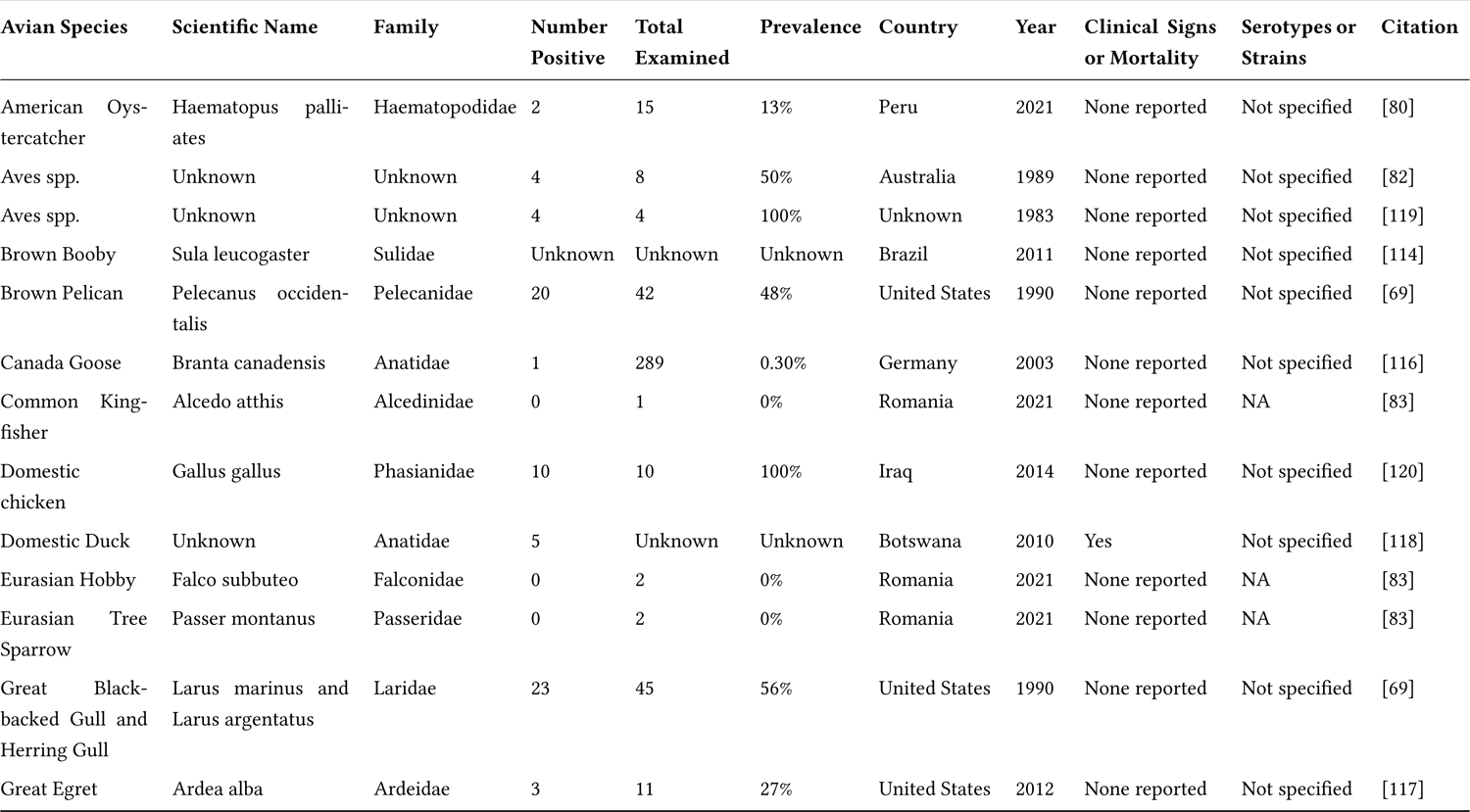
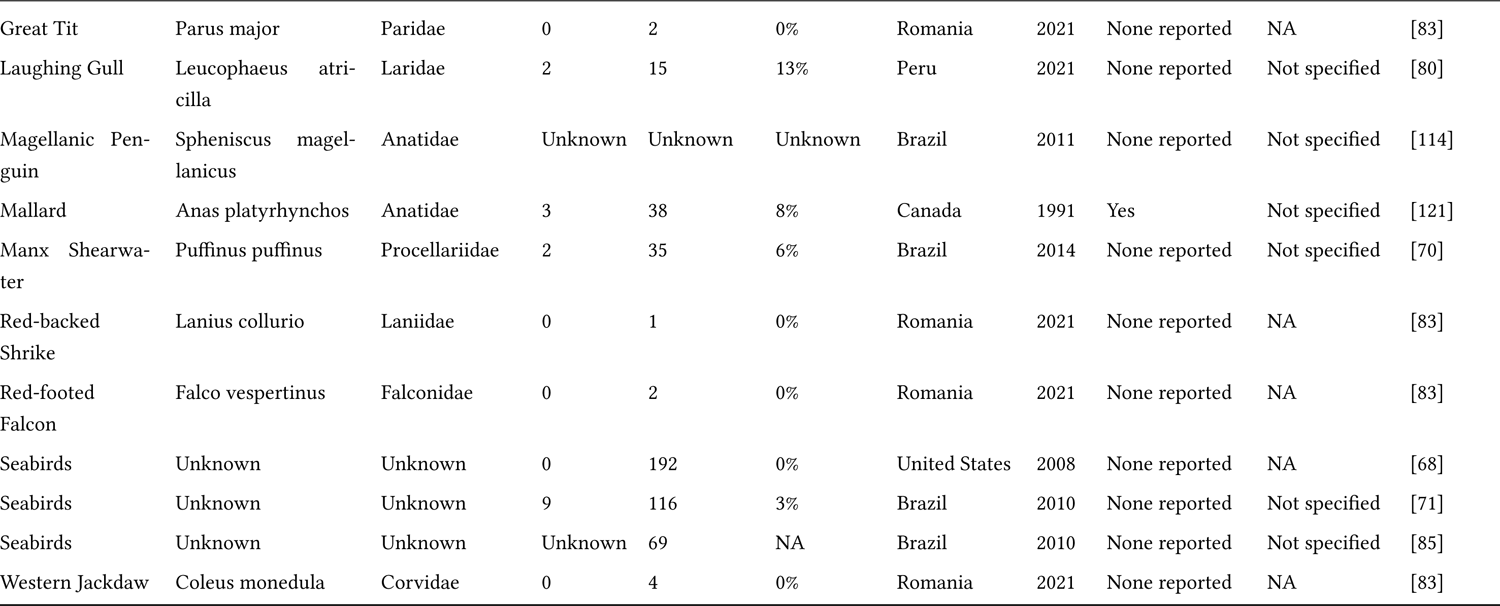
This table details the records extracted from the 15 studies that investigated birds as hosts for *Vibrio alginolyticus*. In the table are provided the common name, the scientific name, the family, the number of birds that tested positive for *Vibrio cholerae*, the total number of birds examined for *Vibrio alginolyticus*, and the prevalence for that record. We also provide the country that the study was performed in, as well as the year the study was conducted or published. For studies that reported clinical signs or mortality, we placed a “Yes” in that column.

#### 3.1.1 Vibrio cholerae

A total of 41 studies in the primary literature examined the role of *Vibrio cholerae* in wild or domestic birds [49, 53, 62–101]. One hundred and fifty-six study records investigated the presence or absence of *Vibrio cholerae*, with the most common technique utilized being culture alone, followed by culture and PCR, or PCR coupled with sequencing. Twenty-five study records reported multiple serotypes from the same species, in the same study. Five of those 25 study records dealt with individual birds who either excreted or displayed multiple serotypes within the same fecal or blood sample or were sampled longitudinally and subsequently cultured positive for different serotypes at different times [53, 91]. Four study records reported the detection of *Vibrio cholerae* O1 from within one or a flock of birds, with Inaba and Ogawa each reported at least once [53, 84, 87, 88]. Serotype distribution across species or taxonomic groups was not analyzed, since the number of birds positive for each serotype was usually not provided in the primary literature. We do, however, report the available data in Table 1. The most common ‘type’ of *Vibrio cholerae* reported from birds was non-O1/O139, however, many study records did not identify or report the serotype of *Vibrio cholerae* that was isolated. *Vibrio cholerae* O139 was not reported from any study.

One-hundred and seven (n = 107) species were examined for the presence of *Vibrio cholerae* antigens or antibodies. An additional sixteen records were extracted from the literature, but we were not able to identify those study records to species. The Anatidae (waterfowl) represented 49 study records, Laridae (gulls and terns) represented 20 study records, and the Ardeidae (shorebirds) represented 10 study records. Within our metaanalysis, 5492 reported birds were tested for *Vibrio cholerae*, and 864 reported birds tested positive, for an overall meta-analysis prevalence of 16%. Study prevalences ranged from 100% in case reports to zero, for example, this often represented rarely captured species that did not yield evidence of exposure to the pathogen. Mallards (*Anas platyrhynchos*) (n = 381) [53, 73, 90, 94, 99] appeared to be the most captured and examined wild species, while domestic chickens (n = 552), both backyard and experimentally inoculated, were the most commonly examined domestic species [65, 76, 87, 88, 91, 97]. Wilson’s Plover (*Charadrius wilsonia*), a species of shorebird that was examined in Venezuela (n = 16/16), had the highest cross-sectional study prevalence for any wild bird captured, with a prevalence of 100% [80]. This was followed by Greater Yellowlegs (*Tringa melanoleuca*), also in Venezuela (n = 6/6), with a prevalence of 100% [80], and Killdeer (*Charadrius vociferus*) in the western United States (n = 13/15), with a prevalence of 86.7% [53].

Clinical signs were reported from 20 study records and were most often associated with *V. cholerae* non-O1/O139 [63, 66, 75, 77, 87, 89, 93, 96, 98, 100, 101]. One study reported clinical signs, primarily edema and cellulitis of the gastrointestinal tract, with an experimental inoculation of O1 Ogawa in domestic chickens [87]. Clinical signs from the literature ranged from respiratory signs to lethargy and sepsis; most infections were associated with other pathogens. However, in a mortality study of American Flamingoes (*Phoenicopterus ruber*), V. cholerae infection was associated with lead toxicity [63]. The largest cross-sectional study to examine wild birds who had exhibited clinical signs in the wild was performed in China, whereby Ruddy Shelducks (*Tadorna ferrginea*) (n = 25/55) tested positive for V. cholerae non-O1 [101]. This study also examined other taxa of birds, such as waterfowl, gulls, shorebirds, and Great Cormorants (*Phalacrocorax carbo*) for the presence of *Vibrio cholerae*, however, study-wide prevalences were generally low when associated with clinical signs (Table 1).

#### 3.1.2 Vibrio parahaemolyticus

We identified 20 studies in the literature that examined the role of wild birds as hosts for *V. parahaemolyticus* [24, 68–73, 82, 85, 95, 102–109]. We extracted seventy-three study records from these papers that examined the prevalence of the pathogen, however, in an additional two study records, we were unable to determine the number of birds infected and/or the number of birds tested [85, 105]. One paper examined the immunogenicity of *V. parahaemolyticus* and *V. vulnificus* in Japanese Quail eggs (*Coturnix coturnix*), and found that birds elicited a high humoral response to the antigens, as measured by ELISA and Western Blots [105]. Most studies utilized culture to determine the presence of *V. parahaemolyticus*, or suckling mice coupled with culture, however, PCR and sequencing was more commonly utilized in more recent works. The Anatidae were represented by 21 study records, the Phasianidae (turkeys, chickens, and pheasants) represented nine study records, and the Laridae represented six study records. We were able to identify 60 species that had been examined for *V. parahaemolyticus*, representing 22 families. For eleven study records, we were unable the species or family of the birds involved in the study [68, 71, 82, 85, 95, 100, 104]. Common Loons (*Gavia immer*) were the most common species tested for *V. parahaemolyticus*, after a multi-year mortality event in Florida [102], however the prevalence was only 0.23% (1/434).

Similar to *V. cholerae*, prevalences for *V. parahaemolyticus* ranged from 100% in the cases of individual study records that were examined, or zero when relatively cryptic and/or scarce species were assessed. Out of the seventy-five study records that we extracted, only 44 reported study records contained birds that tested positive for the pathogen. The highest prevalence for wild birds captured in a cross-sectional study was 68%, involving three species of gulls: Herring Gulls (*Larus argentatus*), Laughing Gulls (*Leucophaeus atricilla*), and Ring-billed Gulls (*Larus delawarensis*) captured off the coast of Florida [69]. This was followed by Herring Gulls and Black-tailed Gulls *(Larus crassirostris*) captured off the coast of Japan, with a prevalence of 67% [106]. Across our study records, we found that 3996 birds had been tested for the presence of the pathogen or for antibodies against the pathogen. A total of 761 birds were positive for *V. parahaemolyticus*, for a meta-analysis prevalence of 19%. Clinical signs were only reported for two studies, in both, co-infection with other organisms was noted [102, 108]. Four studies were associated with other *Vibrio spp.* that were not identified to species [69, 73, 107, 109]. Few studies overlapped between reporting both *V. cholerae* and *V. parahaemolyticus* in birds [69, 73, 85].

#### 3.1.3 Vibrio vulnificus

Eight studies reported examining wild or domestic birds for the presence of *V. vulnificus* or *V. vulnificus* antibodies in the literature, from which we were able to extract 21 study records [71, 83, 85, 105, 106, 109–111]. At least 17 species were represented in this dataset, categorized into 10 families. We were unable to determine within-study prevalences for five of those 21 study records, however, due to the pooling of samples [85, 105, 111]. The most commonly utilized method of determining exposure to *V. vulnificus* in birds was the use of a biochemical panel coupled with culture [83, 110]; similar to *V. parahaemolyticus*, PCR and sequencing were more commonly used in later papers [109, 111]. Study prevalences ranged from zero for rarely captured and/or examined species, to 50%, which was attributed to one of two Black-tailed Godwits (*Limosa limosa*) captured in China that was positive by PCR and sequencing [109]. This was followed by a prevalence of 26% for Herring Gulls and Black-tailed Gulls sampled off the coast of Japan [106]. The largest cross-sectional study was performed in Ogun State, Nigeria, from which multiple farms, representing 565 domestic chickens, were sampled for the presence of exposure to *V. vulnificus* [110]. The study wide prevalence was 0.7%.

No clinical signs or mortality events were reported from any study. In a cross-sectional sampling of urbans birds in Houston, Texas, Zhao et al. 2020 reported [111] that Muscovy Ducks (*Cairina moschata*) and Laughing Gulls excreted more *V. vulnificus* (vvh) than American Crows in the winter as compared to the summer. The greatest diversity of pathogenic *Vibrio* species was reported from a study of stranded seabirds (n = 17/69, *Vibrio spp.*, prevalence of 25%) in Brazil, from which *V. vulnificus* was isolated along with *V. cholerae, V. parahaemolyticus, V.cincinnatiensis, V. fluvialis, V. harveyi*, and *V. mimicus* [85]. The individual prevalence of *V. vulnificus*, or the species afflicted in this study was not reported, however a follow-up study reported a *V. vulnificus* prevalence of 1.7% in Brazilian seabirds [71]. Overall, 1231 birds were examined for evidence of exposure to *V. vulnificus*, and 94 were positive, for a meta-analysis wide prevalence of 8%.

#### 3.1.4 Vibrio alginolyticus

In the literature, we recovered 15 studies in which the role of domestic and wild birds as hosts for *V. alginolyticus* was examined, providing us with 49 study records [68–73, 83, 90, 102, 105, 110, 112–115]. Two study records, involving seabirds off the coast of Brazil, did not identify the sampled birds to species [71, 85]. Forty-seven species were represented in this data subset, categorized into 18 families. Nineteen study records were attributed to the Anatidae, five study records to the Laridae, and three study records were represented by the Falconidae family, known for its small falcons and hawks. Culture, followed by biochemical panels, were the most commonly utilized methods to identify the pathogen. PCR was rarely utilized. The highest prevalence of *V. alginolyticus* recovered from a cross-sectional study of wild birds involved Herring Gulls, Laughing Gulls, and Ring-billed Gulls captured off the coast of Florida, with a prevalence of 68% [69]. The next highest prevalence of the pathogen was 55%, originating from Herring Gulls and Great Black-backed Gulls captured along coastal Connecticut [69]. Across the board, prevalences ranged from zero to 68%, no study record reached a prevalence of 100%. In a study performed in Ogun State, Nigeria, *V. alginolyticus* was isolated from 2% of domestic chickens [110].

Clinical signs and mortality were recorded by two studies, one involving a multi-year mortality event of Common Loons in Florida, and the second involved a mortality event off the coast of Oahu, Hawaii, of Wedge-tailed Shearwaters (*Ardenna pacifica*), which demonstrated prevalence of 10% [102, 115]. Clinical signs ranged from emaciation and lethargy to toxemia and sepsis; bacteremia in the case of the Wedge-tailed Shearwaters was strongly suspected [102, 115]. In a diagnostic examination of critically endangered Mauritius Kestrels (*Falco punctatus*), a captive individual (1/6) was positive by culture for *V. alginolyticus*, yet no clinical signs were noted [113]. In general, waterfowl demonstrated the lowest prevalences for any group of birds, besides passerines, for the pathogen [73, 83]. Throughout our dataset, 258 birds of 2,967 sampled birds tested positive for *V. alginolyticus*, for a meta-analysis prevalence of 9%.

#### 3.1.5 Vibrio fluvialis

From the literature, we found 15 studies that reported examining wild or domestic birds for the presence of *V. fluvialis* or *V. fluvialis antibodies*, from which we were able to extract 26 study records [68, 69, 71, 71, 80, 82, 83, 85, 105, 114, 116–121]. At least 20 species were reported in this dataset, representing 15 families. Four records did not provide sufficient data from which to identify birds to species or family. Culture was the most commonly utilized method to identify *V. fluvialis*, followed by a biochemistry panel. The largest cross-sectional study examining the prevalence of *V. fluvialis* in wild birds was performed on Canada Geese in Germany, however, only one of 289 birds cultured positive for the pathogen [116]. *V. fluvialis* was associated with one mortality event – a die-off of overwintering Mallards in Canada which was attributed to a Vitamin A deficiency [121]. The reported prevalence of the pathogen for these birds was 8%. In a study of captive study of Great Egrets (*Ardea alba*) captured from the Mississippi Delta, control birds shed *V. fluvialis* for four of seven days in captivity [117]. The highest prevalence was associated with a study performed in Connecticut involving Herring Gulls and Great Black-backed Gulls, with 55% culturing positive for *V. fluvialis*. Studies with a prevalence of 100% involved two experiments, one involving of avian-sourced strains, and a mitogenicity study on domestic chickens [119, 120]. Overall, the meta-analysis prevalence, including experimental infection studies (88/834) was 11% percent (85/834).

#### 3.1.6 *Other Pathogenic* Vibrio spp.: V. cincinnatiensis, V. hollinsae, e.g., Grimontia hollinsae, V. furnissii, V. mimicus, V. harveyi, V. scophthalmi, V. metschnikovi*, and* Photobacterium damselae

The abundance of studies that reported on other pathogenic *Vibrio* species that were isolated from wild or domestic birds varied (Table 6). *V. cincinnatiensis* was examined by five studies and provided five study records [42, 70, 71, 85, 114]. Unspecified seabirds were the taxa that were examined most frequently [71, 85], however overall prevalences were low across all studies for a mean prevalence of 3%. The presence or absence of *Photobacterium damselae* was examined by three studies [69, 102, 122], and yielded three study records from the United Kingdom and the United States. Two studies involved mortality events, one of Common Loons in Florida, and the second of British passerines [102, 122]. Across these cross-sectional studies, the overall meta-analysis prevalence was approximately 5%. Our literature search of *V. furnissii* yielded four study records from three cross-sectional studies [71, 81, 114], two of those study records did not identify the number of birds positive or the number of individuals examined. The Laridae were the most prevalent species identified in association with *V. furnissii*, specifically Kelp Gulls (*Larus dominicanus*), Laughing Gulls, and Brown Boobys (*Sula leucogaster*). The overall meta-analysis prevalence for this *Vibrio* pathogen was approximately 2%.

**Table 6:**
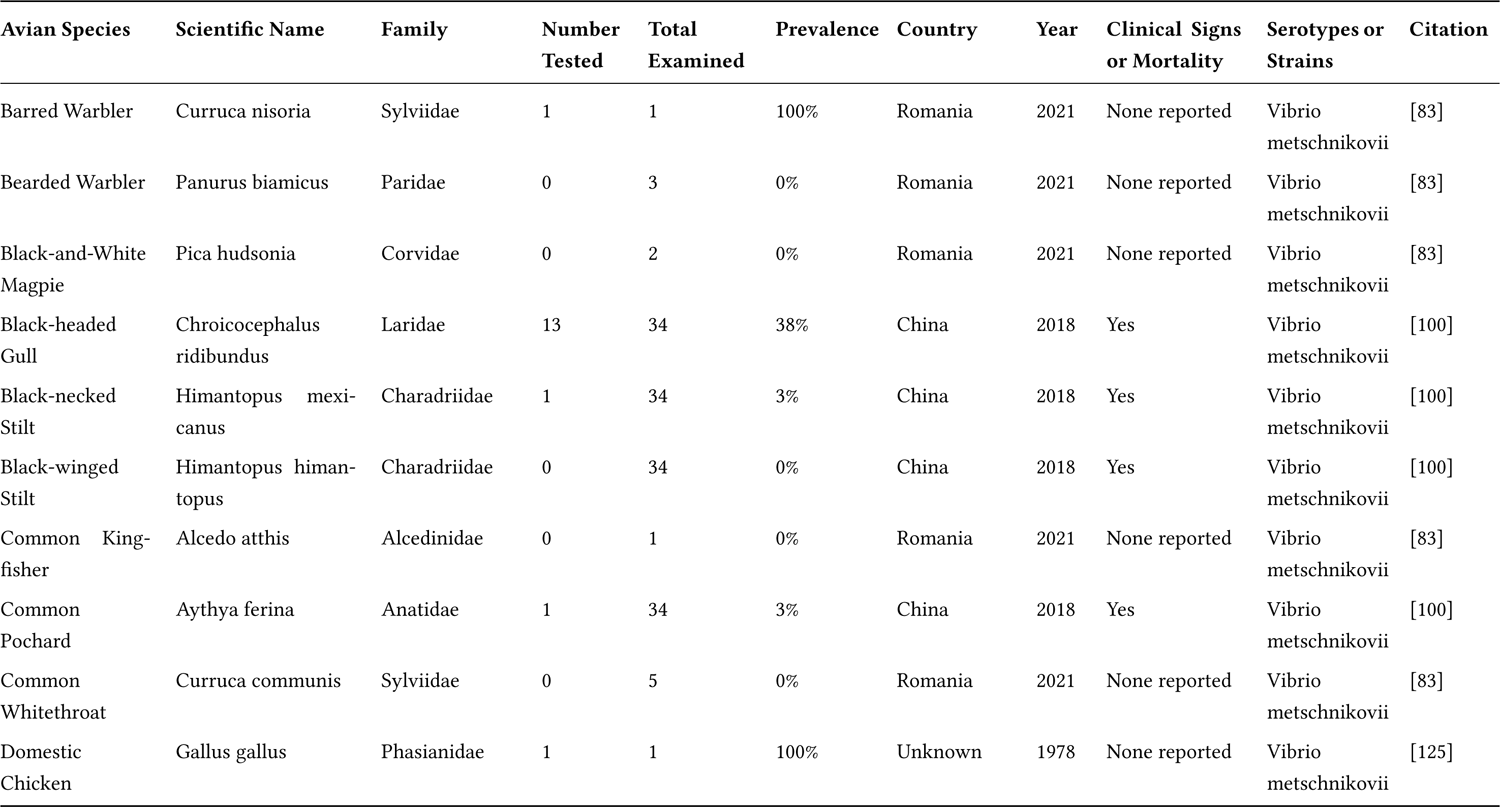
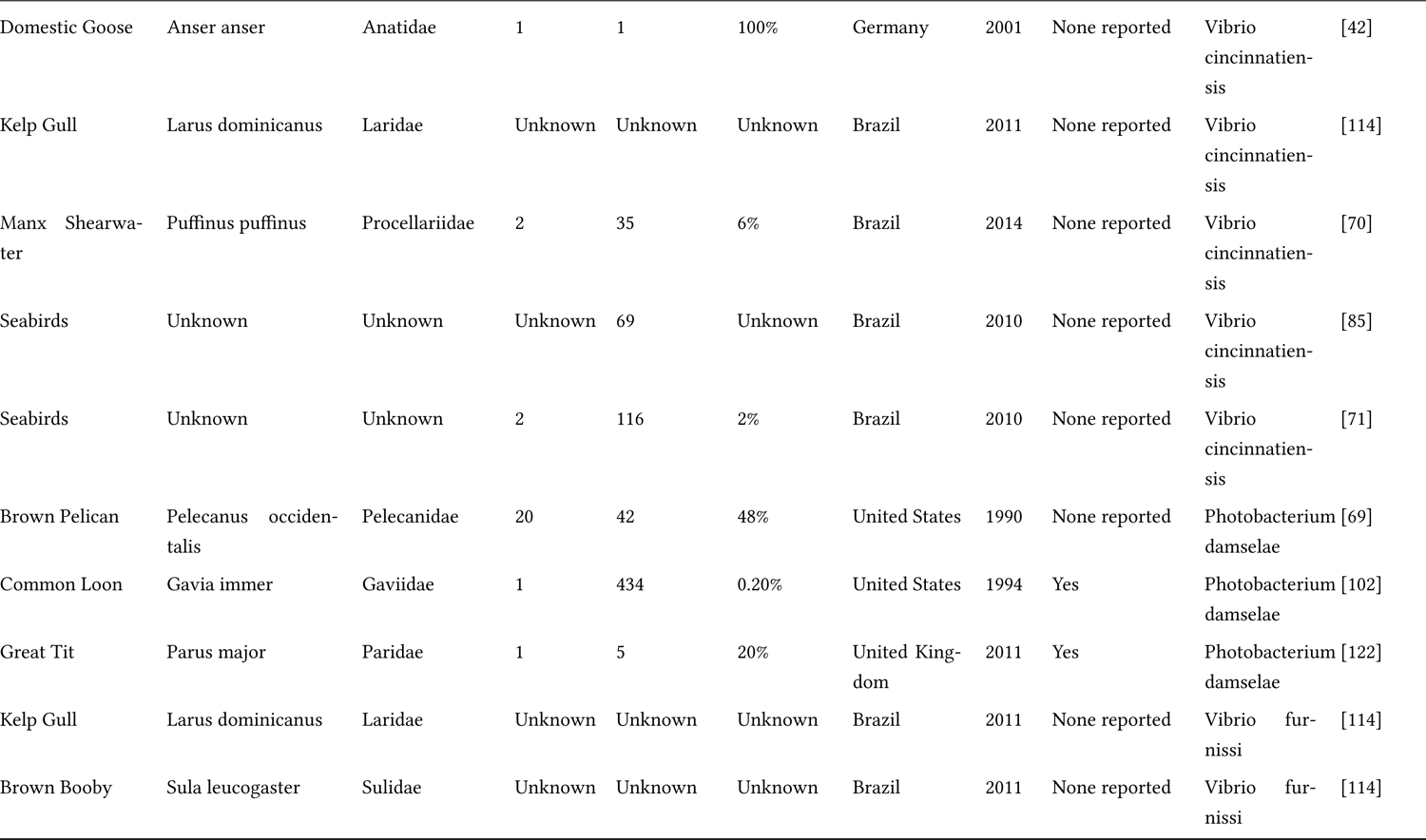
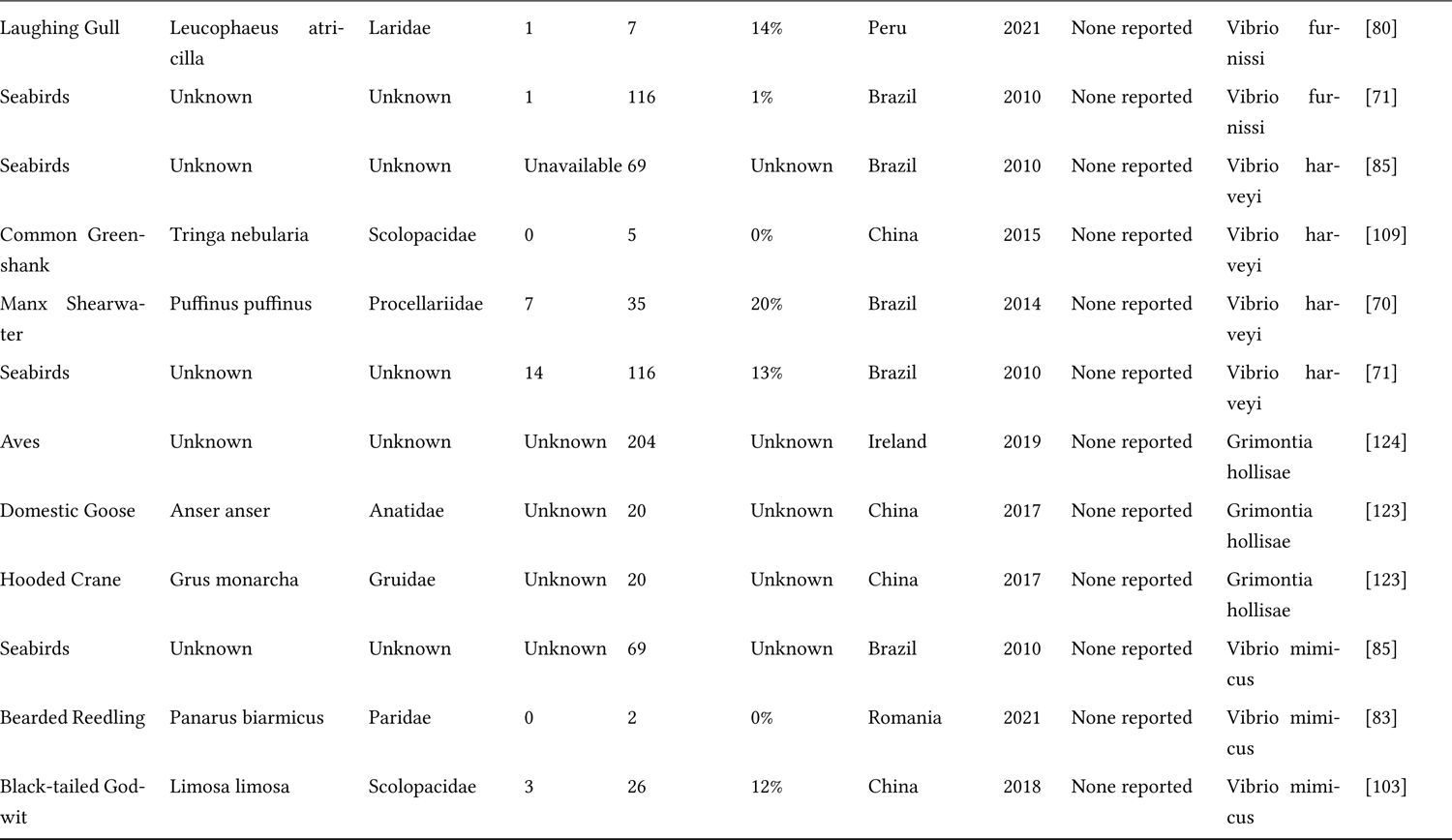
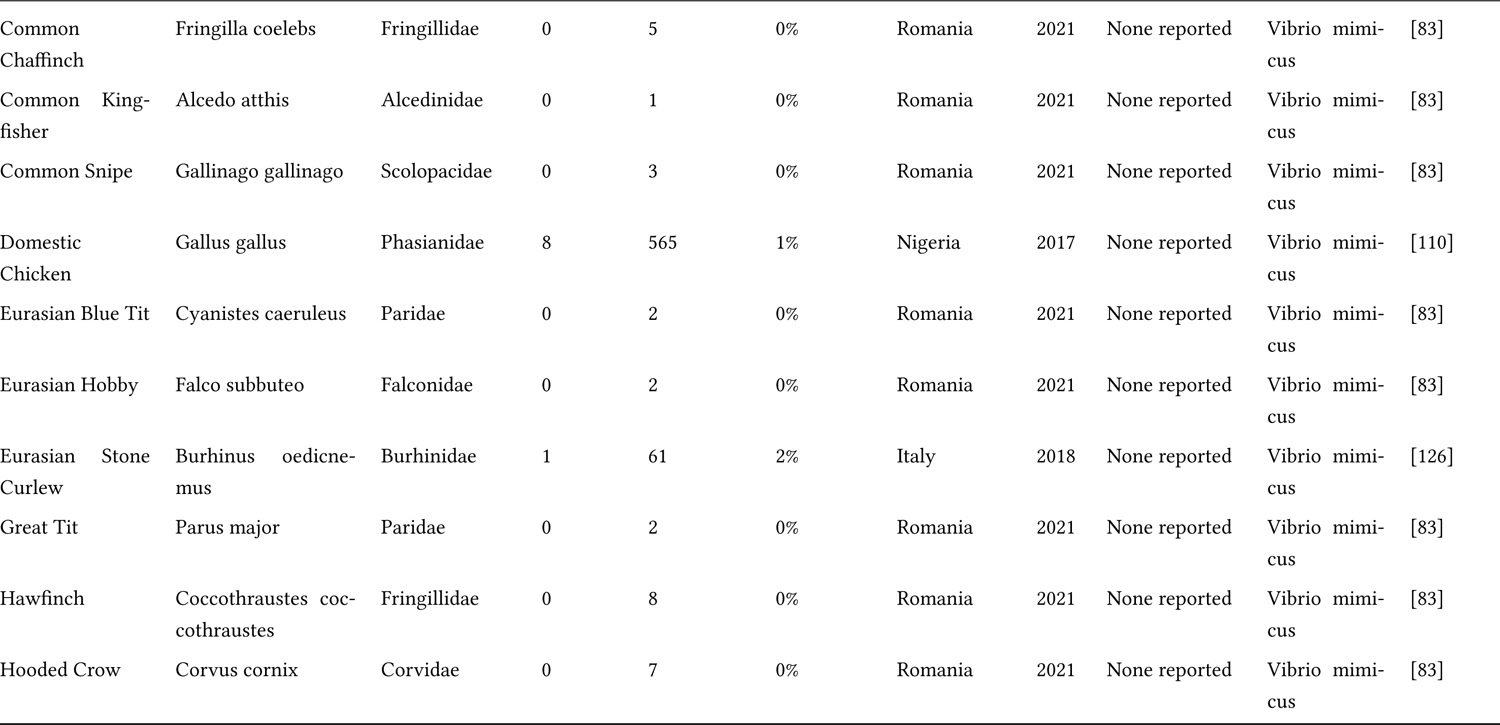
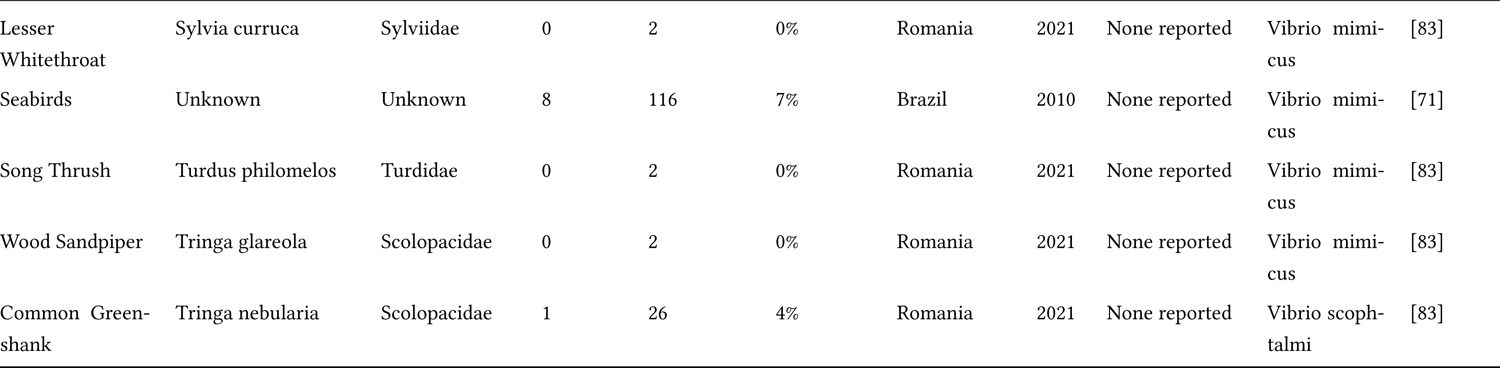
This table details the records extracted from the 18 studies that investigated birds as hosts for other pathogenic *Vibrios*, including *V.metschnikovii, V. mimicus, V. cincinnatiensis, V. scophtalmi, Grimontia hollisae*, formerly *V. hollisae, Photobacterium damselae*, formerly *V. damselae, V. furnissii*, and *V. harveyi*. In the table are provided the common name, the scientific name, the family, the number of birds that tested positive for the respective pathogen, the total number of birds examined, and the prevalence for that record. We also provide the country that the study was performed in, as well as the year the study was conducted or published. For studies that reported clinical signs or mortality, we placed a “Yes” in that column.

The search for studies involving *V. harveyi* and avian species yielded four cross-sectional studies and four study records [70, 71, 85, 109], involving seabirds and Manx Shearwaters (*Puffinus puffinus*). A single study involving a coastal sandpiper, the Common Greenshank (*Tringa nebularia*), had a prevalence of 0% out of five birds that were tested by PCR [109]. From the two studies that provided individual birds positive in contrast to individual birds examined, we were able to calculate a *V. harveyi* prevalence of approximately 13%. No study that tested for this pathogen reported clinical signs or a mortality event. *Grimontia hollisae* was rarely detected in birds, as we found only two studies, resulting in three study records, that searched for the pathogen in avian hosts [123, 124]. One study examined the shared microbiota between wild Hooded Cranes (Grus monacha) and domestic geese (*Anser anser*) using Miseq – *Grimontia hollisae* was identified as a potential pathogen, but the total number of birds colonized was not reported [123]. A longitudinal microbiome study involving shorebirds off the coast of Cork, Ireland discovered *Grimontia hollisae* in fecal samples, however, the number of samples positive/examined was not enumerated [124]. Clinical signs or mortality were not reported from either study.

*Vibrio metschnikovii* was reported from three studies, resulting in 10 study records [83, 101, 125]. Approximately half the study records examined passerines of Romania as hosts [83], including members of the Laniidae, Sylviidae, and Paridae families, all of which were negative for the pathogen by biochemical panels. The highest prevalence was reported from sites in Inner Mongolia, China, by Black-headed Gulls (*Chroicocephalus ridibundus*), from which an overall study prevalence of 38% was reported [101]. Clinical signs and a mortality event that spanned multiple waterfowl and waterbird species were documented in the study by Zheng et al. 2021 [101]. The number of total tested birds were not available for analysis in the latter study; thus, we could not report a meta-analysis prevalence of *V. metschnikovi* with confidence. *Vibrio mimicus* was examined by six studies, yielding 18 study records [71, 83, 85, 110, 126]. Biochemical panels were the most common diagnostic tool used to identify the pathogen, however, overall prevalences were very low across studies. In a study of wading birds and songbirds performed along the Danube Delta of Romania, all sampled birds (n = 38) were negative for *V. mimicus* [83]. On the other hand, wading birds and seabirds sampled in China, Brazil, and Italy demonstrated evidence of shedding the pathogen [71, 103, 126]. The only study to examine the role of domestic birds as hosts was performed in Ogun State, Nigeria – this study yielded a prevalence of approximately 1% [101]. Across studies, 806 birds were examined for the presence of the pathogen, with 21 testing positive, resulting in a meta-analysis prevalence of 2%. No clinical signs or mortality events were reported from any study that examined the role of birds as hosts for *V. mimicus*.

*Vibrio scophthalmi* was only reported from one study, resulting in a single study record [103]. A Common Greenshank (1/26) that was sampled using whole genome sequencing was positive for the pathogen [103]. This study was not associated with clinical signs or a mortality event. Uncategorized *Vibrio* spp. were reported from 14 studies [68, 69, 71, 73, 74, 80, 83, 95, 100, 109, 124, 127–129]. Two studies were associated with clinical signs and/or mortality events, however, these outbreaks were attributed to other causal pathogens [100, 127]. Culture, followed by biochemical panels, were the most commonly utilized methods of identifying *Vibrio* spp. Given that many studies did not identify these *Vibrio* spp. to species or identify the number of birds excreting them, we were unable to calculate a meta-analysis wide prevalence.

## 4 Discussion

The question of pathogenic *Vibrio* spp. as the etiological agents of disease in birds remains only partially answered. Of the 76 studies that surveyed birds for pathogenic *Vibrio* species, 19 reported disease or death from individuals, scaling up to community level events [63, 66, 75, 77, 87, 89, 93, 96, 98, 100, 101, 108, 115, 118, 121, 125, 127]. Yet, it remains uncertain whether these pathogenic *Vibrio* species are opportunistic pathogens that contribute to morbidity and/or mortality in already stressed individuals, or whether they can be the primary arbiters of disease [111]. Experimental inoculation studies reported contrasting results, if they reported clinical signs at all [49, 87, 99, 120]. In addition, avian susceptibility to pathogenic *Vibrio* species may also be conflated by host species, natural history, and prior exposure, resulting in an as yet-understood degree of immunity [130, 131]. In our meta-analysis, disease was most commonly associated with *V. cholerae*, followed by *V. metschnikovi* and *V. parahaemolyticus* – notably, 11 of 39 study records were associated with domestic ducks (*Anas platyrhynchos* or *Anser anser*) or domestic chickens [66, 67, 75, 77, 87, 96]. This may have implications for agriculturally associated species in areas of the world where backyard birds are the primary protein source for pastoral families [132–134].

Of the 425 study records we extracted from the literature, interestingly, the Anatidae represented 105 of them, including wild and domesticated Mallards, which represented 16 study records. The Laridae represented 39 study records, prominently represented by Laughing Gulls, Herring Gulls, and Ring-billed Gulls. Shorebirds and waders, categorized into the Ardeidae family, represented 16 study records, primarily of egrets and herons. These bird species are often highly associated with coastal estuarine and marine environments [135–137], which are also inhabited by autochthonous and halophilic *Vibrio* species. These results are congruent with what is known of avian foraging ecology and *Vibrio* habitat specificity [6, 36, 55, 138–140]. What was unexpected were the number of ground-foraging birds that tested positive for pathogenic *Vibrio* species that are often not strictly associated with aquatic environments, such as Great Tits (*Parus major*), Garden Warblers (*Sylvia borin*), and Hooded Crows (*Corvus cornix*) [81, 83].

For example, in a study of Egyptian backyard poultry (chickens, turkeys, and waterfowl), 36% of examined birds were positive for*V. cholerae*, including chickens and turkeys [76]. Domestic chickens accounted for 13 total study records, across geographical areas as varied as the United States, Bangladesh, Egypt, Ghana, Nigeria, Iraq, and India, and reported as early as 1972 [65, 76, 87, 88, 91, 97, 110, 112, 120, 125]. On the other hand, another surprising result was the low prevalence of pathogenic *Vibrio* species cultured from seabirds that were sampled from the New England region of the United States, with only one of 192 birds testing positive for *Vibrio cholerae*, non-O1 [68]. This result may be due to several reasons, many of which are not mutually exclusive [137]. For one, as seabirds tend to spend more time in marine versus coastal habitats, they may be less susceptible to exposure from pathogenic *Vibrio* species that tend to congregate in lower salinity, brackish habitats [141]. In addition, the northern Atlantic may harbor a lower abundance of pathogenic *Vibrios* during the cooler months as a result of low sea surface temperatures [13] [12, 142]. Lastly, it may be possible that although pathogenic *Vibrio* spp. may cause disease in seabirds, that the recovery of carcasses or diseased individuals may be reduced due to minimal mortality, low carcass persistence, and increased distances from urbanized centers [143–145].

Meta-analysis prevalences varied across pathogenic *Vibrio* species, but all were below 20% (e.g., 19% for *V. parahaemolyticus*, 16% for *V. cholerae*, 13% for *V. harveyi*, 11% for *V. fluvialis*, 9% for *V. alginolyticus*, 8% for *V. vulnificus*, 5% for *P. damselae*, 2% for *V. furnissi*, and 1% for *V. mimicus*). Given that we utilized experimental inoculation studies coupled with cross-sectional studies, there is likely a degree of reporting bias in our meta-analysis prevalences [146], however, we speculate that this reporting bias is likely offset by the reportedly few studies that have targeted these pathogens for investigation in wild and domestic birds. To determine true ‘prevalence’, and avian susceptibility under ecological conditions, longitudinal studies that sought to recover these pathogens from a community of birds would be more informative [147, 148]. In addition, these studies would need to utilize large sample sizes, as well as represent various ecological foraging guilds, in geographic locations with both low and high recovery rates of these pathogens from their aquatic, environmental reservoir [149–152].

With a meta-analysis *Vibrio* prevalence of 16% coupled with the reports of clinical signs, there is a possibility that pathogenic *Vibrio* species—specifically *V. parahaemolyticus*, *V. cholerae*, and *V. metschnikovi*—may be emerging pathogens of wild and domestic aquatic or wetland birds [153, 154]. Gire et al. 2012 [155] defined emerging pathogens as falling into two categories: introduced microbes, and existing microbes that rapidly increase in prevalence and/or incidence in a population. Given that so little is known of non-cholera *Vibrio* species in human hosts, however, it is difficult to distinguish between the two categories in our avian hosts given the currently available data. Speculation suggests that these *Vibrio* pathogens may have a long-standing relationship with aquatic birds. However, as climate change alters and influences the abundance and distribution of pathogenic *Vibrio* species in marine and estuarine environments, so too may the incidence of these pathogens in wild and domestic birds [156].

In summary, we have offered a rigorous meta-analysis that examines the prevalences of *Vibrio* spp. across bird species. In doing so, we reveal both a plethora of data that fortify the notion that birds are an underappreciated object of study and are potential reservoirs for pathogenic bacterial species. In the context of a dynamic ecology defined by climate change, and human-associated activities, we suggest that bird reservoirs should be the focus of more rigorous study, as they may be an actor in *Vibrio* emergence events. Transcending the case of birds, our species propose that more attention should be paid to other animal species that may harbor pathogens of interest to human health.

